# A High-Throughput Biocatalytic Platform for Screening Isomeric Kainoid Natural Products

**DOI:** 10.1101/2025.09.07.674781

**Authors:** Robert A. Shepherd, Manasa Ramachandra, Austin R. Hopiavuori, Melanie C. Jones, Conrad A. Fihn, Alex J. Tabag, Ananya Manjunath, Limar Y. Gad, Chloe R. Whipple, Shaun M. K. McKinnie, Laura M. Sanchez

**Affiliations:** Department of Chemistry and Biochemistry, University of California Santa Cruz, Santa Cruz, CA 95064

**Author notes:** These authors contributed equally to this work (Robert A. Shepherd, Manasa Ramachandra).

## Abstract

Random mutagenesis generates large, diverse enzyme libraries, however high-throughput screening is challenging when transformations yield isomeric small molecule products. We developed a matrix-assisted laser desorption/ionization–trapped ion mobility spectrometry–mass spectrometry (MALDI-TIMS-MS) platform for rapid, label-free screening of libraries of kainoid synthases, Fe/αKG-dependent cyclases that produce the isomeric neurotransmitters kainic acid (KA) and kainic acid lactone (KAL). This platform achieves near-baseline separation of KA and KAL directly from microbial biomass, with parallel reaction monitoring–PASEF (PRM-PASEF) enhancing signal-to-noise for isomeric product ratio analysis. Using this workflow, we screened a 318-variant DsKabC/GfKabC DNA shuffle library in under 30 minutes, identifying seven variants with improved KAL conversion. Two variants showed a near-complete shift toward KAL production with greater substrate consumption, while retaining favorable expression profiles compared to GfKabC. These results establish MALDI-TIMS-MS as a generalizable platform for high-throughput isomeric product screening and provides new kainoid synthase variants to probe structure–function relationships for further development as biocatalysts. Importantly, this screening approach can be broadly applied to interrogate enzyme variants acting on small molecules, delivering rapid and accurate quantitative insights without reliance on chromatography, enzyme purification, or substrate derivatization.

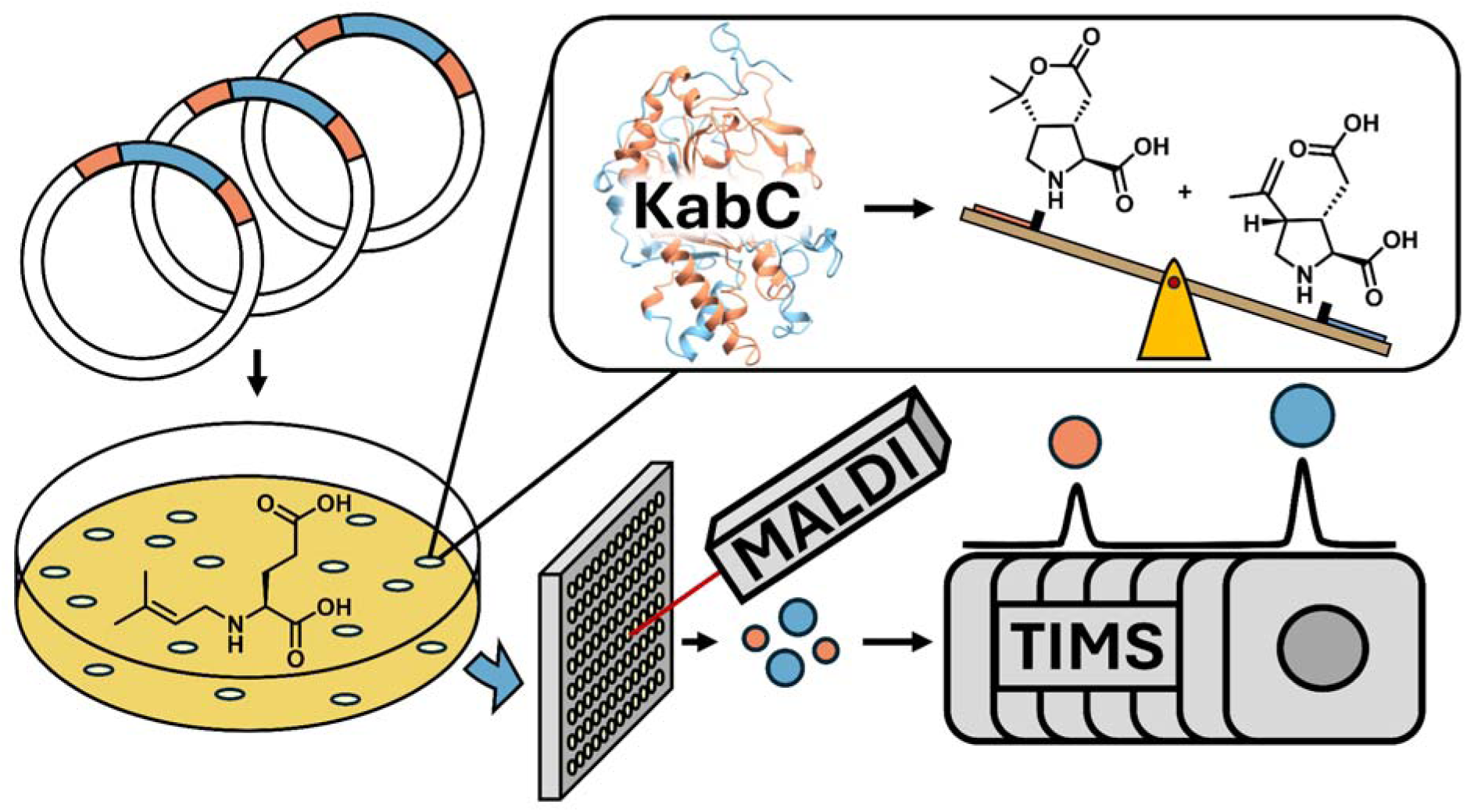

## Introduction

Random mutagenesis techniques have significantly enhanced our ability to rapidly generate large libraries with unprecedented genetic diversity.^1,2^ For example, directed evolution (DE) of heme proteins has produced enzymes capable of catalyzing complex, *new-to-nature* chemistries such as C–H amination and the formation of chiral organoboranes.^3,4^ However, random mutagenic libraries often contain hundreds to thousands of variants, making the identification of those variants capable of performing highly specialized chemical reactions a major bottleneck. Notable progress has been made using automated screening and selection technologies—such as microfluidic sorting,^5,6^ digital imaging,^7,8^ and microtiter plate workflows^9–13^—paired with optical methods to enable high-throughput enzymatic screening. While these approaches effectively address throughput challenges, many depend on absorbent, fluorescent, or chromogenic components and require optimization for specific biochemical reactions, which limits their general applicability.

Mass spectrometry (MS)-based reaction screening approaches have gained traction over the past decade due to their label-free nature, making them highly adaptable to a wide range of screening scenarios.^14^ While MS generally sacrifices throughput, its broad applicability is a major advantage. However, MS is only suitable for transformations in which the substrate and/or products exhibit a detectable difference in mass. This limits the utility in biochemical systems that generate multiple isomeric or isobaric products, as MS alone may not have the resolution to distinguish between them. Orthogonal methods such as liquid chromatography (LC) are often employed to resolve isomeric or isobaric analytes prior to MS analysis, further reducing throughput. To address this challenge, we developed a chromatography-free high-throughput workflow to monitor isomeric products using the kainoid synthase family of enzymes as a model system.

Kainoid synthases belong to the broader superfamily of nonheme iron/α-ketoglutarate-dependent dioxygenases (Fe/αKGs), a large and versatile group of enzymes involved in both primary and secondary (specialized) metabolism.^15–18^ Fe/αKGs utilize a mononuclear Fe(II) cofactor, α-ketoglutarate, and molecular oxygen to catalyze oxidative transformations via a radical mechanism.^16–18^ These enzymes support a broad range of chemistries—including hydroxylation, halogenation, ring closure, epimerization, and epoxidation—making them ideal for diverse biocatalytic applications.^19–22^ A small subset of Fe/αKGs function as oxidative cyclases, forming new C–N,^23^ or C–C bonds to construct ring systems. Isolated from various marine micro- and macroalgae, kainoid synthases catalyze the cyclization of various *N*-prenylated L-glutamic acid substrates to generate the pyrrolidine ring characteristic of “kainoid” marine neurotoxins.^18,24–27^ Among them, DsKabC, from the red alga *Digenea simplex*, oxidatively cyclizes its prekainic acid (PKA, *N*-dimethylallyl-L-glutamic acid) substrate to kainic acid (KA)—a potent ionotropic glutamate receptor (iGluR) agonist used extensively in neurological research and murine epilepsy models.^28–30^ DsKabC produces a minor secondary product, kainic acid lactone (KAL) (Figure 1A, C), a bicyclic isomer with antagonistic iGluR activity.^31^ Intriguingly, GfKabC, an ortholog from the red alga *Grateloupia filicina*, produces primarily KAL *in vitro*, with KA being its minor secondary product (Figure 1C).^26^ Although DsKabC and GfKabC share high sequence homology, accept identical PKA substrates, and exhibit highly similar predicted substrate pocket compositions (Figure 1B), they display distinct product profiles (Figure 1C). The mechanistic branch point between KA and KAL formation likely occurs after pyrrolidine ring closure (Figure 1A), however the structural basis underlying this divergence remains unclear.^26,32^

**Figure 1.**
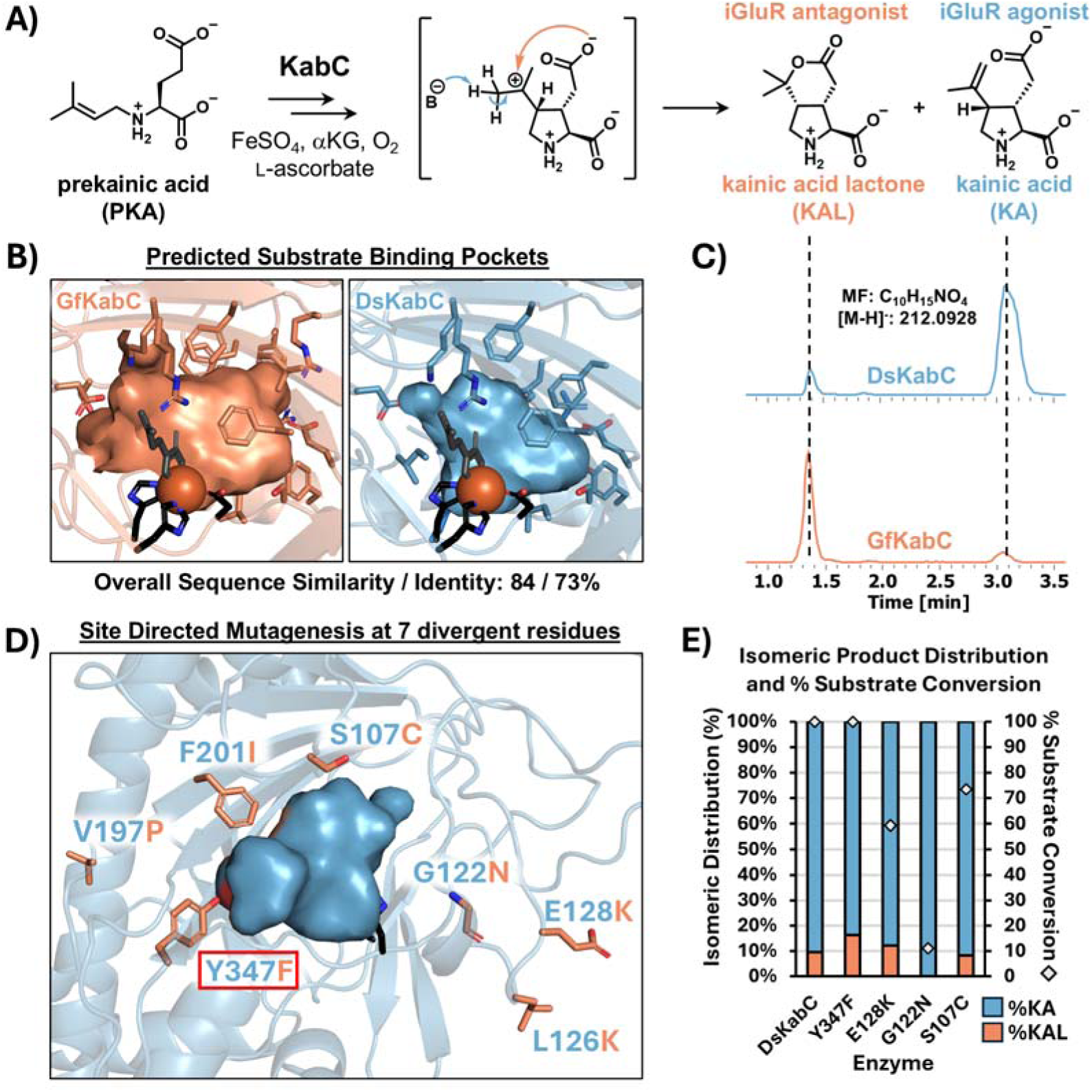
Overview of KabC structure and activity. A) Abbreviated mechanism for the conversion of prekainic acid to both kainic acid (KA) and kainic acid lactone (KAL) by KabC. The mechanistic branch point putatively occurs after pyrrolidine ring formation, with deprotonation resulting in KA formation, and nucleophilic addition of the side chain carboxylate resulting in the formation of KAL. B) The predicted substrate binding pockets of DsKabC and GfKabC are highly similar in composition, reflecting their overall high sequence similarity and identity (84 and 73%, respectively). C) LC-MS extracted ion chromatograms of *m/z* 212.09 ± 0.2, demonstrating the differing production of KA and KAL between DsKabC and GfKabC. D) Seven sites chosen for rational SDM based on multiple sequence alignment of KabC orthologs. Boxed in red is the Y347F mutant carried forward as an additional positive control for screening development. E) *In vitro* isomeric distribution and percent substrate conversion (represented by the gray diamonds) for DsKabC and the four prioritized single-point KabC variants via LC-MS (n = 1). The displayed LC-MS data were collected on a Bruker AmaZon SL.

Screening KabC products at scale, however, presents significant challenges. KA and KAL are constitutional isomers that share the same chemical formula (C_10_H_15_NO_4_) but differ in structure (Figure 1A, C), making them indistinguishable by MS alone. Furthermore, their lack of distinguishable chromophores renders traditional absorption-based enzymatic screening methods ineffective for monitoring their relative abundance. To address this, LC-MS is typically employed as mentioned above. While LC-MS is reliable for quantifying KA and KAL, its lengthy acquisition time—requiring up to 15 minutes per sample—makes it impractical for large-scale screening. We developed a matrix assisted laser desorption/ionization mass spectrometry (MALDI-MS) reaction screening platform due to its high-throughput, label-free nature, enabling the rapid analysis of a diverse range of chemical products.^33–38^ This platform was enhanced by the use of ion mobility spectrometry (IMS) which is a timescale-compatible method for gas-phase ion separation.^39–44^ A unique advantage of IMS is its ability to rapidly separate isomeric species based on their size and 3-D shape.^45–47^ Here, we combine the simple sample preparation and rapid analysis of MALDI-MS with the isomeric separation capability of trapped ion mobility spectrometry (TIMS) to screen a mutant library of KabC enzymes for relative KA and KAL production. Using this MALDI-TIMS-MS platform, we evaluated a total of 1,054 random KabC variants with a total acquisition time of ∼1.5 h. The results showed that random mutagenesis informed by the evolutionary similarity and functional diversity of *dskabC* and *gfkabC* is a powerful approach for rapidly probing how subtle sequence changes—particularly in second-shell or distal regions—influence active site geometry and control isomeric selectivity.

## Results and Discussion

### Semi-rational design through site directed mutagenesis and fusion enzyme construction

Given the absence of kainoid synthase crystal structures, we initiated our studies using AlphaFold 2^48^ to model both enzymes and select specific residues for mutation. Sequence homology across KabC orthologs highlighted residues that are generally conserved but differ between DsKabC and GfKabC. Orthologs from *Palmaria palmata* (PpKabC) and *Rhodophysema elegans* (ReKabC) also produce both KA and KAL, though with lower conversion and a consistent bias toward KA.^26^ Multiple sequence alignment further highlighted thirteen residues conserved in PpKabC, ReKabC, and uncharacterized PhKabC (*Palmaria hecatensis*). In contrast, DsKabC and GfKabC diverge at these positions relative to each other and to the broader ortholog set, suggesting these residues may shape kainoid isomer distribution (Figure S1). Given the reported solubility and stability constraints of GfKabC,^24,26^ the corresponding residues were introduced into the more robust DsKabC scaffold. Seven of the thirteen planned site-directed mutants (SDM) were successfully generated and confirmed by Sanger sequencing (Figure 1D).

DsKabC SDM constructs were initially evaluated using whole-cell lysate reactions and monitored for relative product distribution via LC-MS (Figure S2). Four mutants (S107C, G122N, E128K, Y347F) were prioritized for protein purification (Figure S3) and *in vitro* characterization based on either improved total conversion or increased relative KAL production (Figure S4). Of these, only E128K and Y347F showed measurable increases in KAL abundance—3% and 7%, respectively—relative to wild-type DsKabC (Figures 1E, S4). Counterintuitively, the G122N mutation eliminated KAL production and drastically reduced substrate conversion (Figure 1E). Notably, Y347F was the only mutant to retain the >99% substrate conversion seen in WT DsKabC. Prior studies in similar Fe/αKG-dependent enzymes have shown that even single mutations within the binding pocket can significantly impact activity due to the stringent geometric constraints required for radical-based catalysis.^49^ In the case of DsKabC, however, only modest changes in isomeric selectivity were observed from rational mutagenesis to their GfKabC counterparts.

Given the minimal changes in isomeric distribution observed from DsKabC SDM, we deliberately replaced larger regions with the corresponding GfKabC sequences. These “fusion” enzyme constructs were generated by targeting two regions with low sequence homology: a C-terminal alpha-helical region; and an internal non-flexible loop, and replacing the GfKabC sequence into the DsKabC scaffold (Figure S5). Following *in vitro* reactions with purified fusion KabC constructs, no substantial changes in the relative abundance of KA and KAL were observed, and substrate conversion was greatly reduced (Figures S6, S7). This outcome underscored the complexity of kainoid synthase structure-activity relationships and motivated us to pursue random mutagenesis approaches. Given the throughput limitations of LC-MS for separating regioisomers, we prioritized developing a higher-throughput platform to enable screening of larger mutagenic KabC libraries.

### Optimization of MALDI-TIMS-MS for Kainoid Natural Products

TIMS works by “trapping” ions through a balance between the drag force from a forward-flowing gas and an opposing electric field gradient. Ions are sequentially released by “ramping down” the electric field, allowing separation based on ion mobility (K), which is often reported as the inverse reduced mobility (1/K_0_) for TIMS.^50^ A key feature for our platform development was optimizing the resolution of the peaks (Figure 2A). Resolution in TIMS is primarily governed by the ramp rate—the rate at which the electric field is decreased during ion release—which is determined by both the ramp time (the duration over which the electric field is reduced) and the width of the 1/K_0_ range being scanned.^51,52^ Increasing the ramp time allows ions with similar mobilities more time to separate in the electric field. Likewise, narrowing the 1/K_0_ range while keeping the ramp time constant effectively reduces the ramp rate, enhancing resolution by generating a shallower electric field gradient over a narrow mobility window. For high-throughput screening applications, these parameters can be balanced to optimize both resolution and acquisition speed.

**Figure 2.**
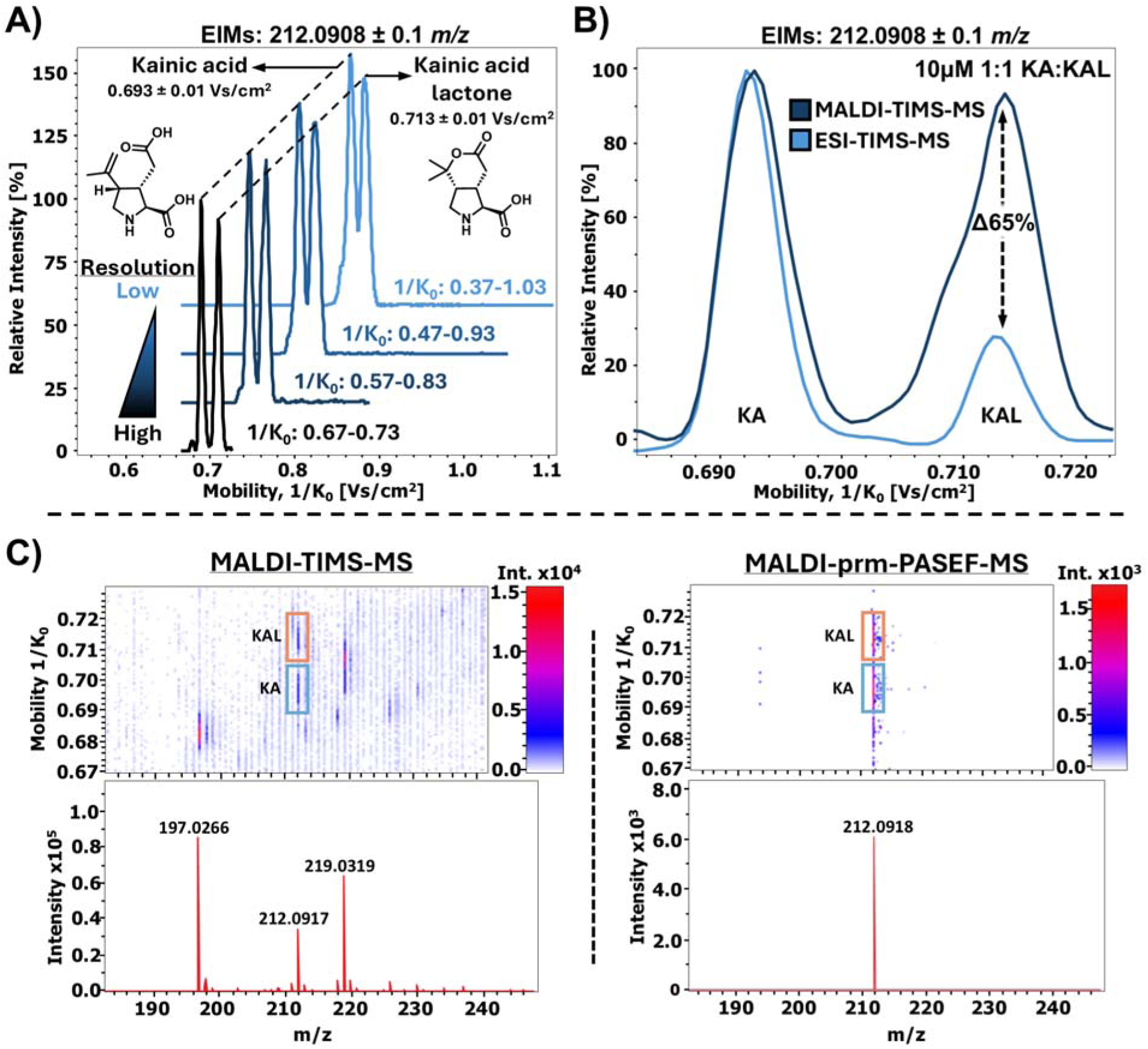
Optimization of TIMS-MS parameters for monitoring KA and KAL. A) Optimization of TIMS resolution for KA and KAL. Increasing the ramp time and decreasing the 1/K_0_ range of acquisition maximized resolution for our analysis. B) Comparison between MALDI- and ESI-TIMS-MS for measuring the relative abundance of KA and KAL. When measuring the same mixture of standards, it was observed that ESI-TIMS-MS displayed a 65% decrease in the relative intensity of KAL, suggesting that MALDI-TIMS-MS is more reliable for relative kainoid quantitation. C) Employment of PRM-PASEF-MS significantly reduced chemical noise for our MALDI-based analysis, simplifying and increasing confidence in downstream data visualization.

To optimize TIMS parameters in real time, we performed direct infusion electrospray ionization (DI-ESI)-TIMS-MS on a 1:1 mixture of pure 10 µM KA and KAL across *m/z* 50– 700 in negative ion mode. Given their structural similarity, we hypothesized that KA and KAL would have nearly identical ion mobilities, supported by predicted collisional cross section (CCS) values of 148.3 and 147.4 Å^2^, respectively.^53^ We initially monitored a broad 1/K_0_ range (0.3–1.5 Vs/cm²) to identify the mobility of *m/z* 212.09. Under these conditions, the mobilogram showed only a single peak, indicating insufficient TIMS resolution to separate the two isomers. Increasing the ramp time from 50 ms to 100 ms revealed a shoulder peak indicative of partial resolution. Extending the ramp time further to 200 ms enhanced separation, making the second peak distinct; however, substantial overlap remained, limiting reliable relative quantification.

Since our analysis was highly targeted for KA and KAL, we opted to reduce the 1/K_0_ range to further enhance resolution and suppress chemical noise. By narrowing the 1/K_0_ range in 20 Vs/cm² increments centered between the peaks of KA and KAL, we observed a substantial improvement in TIMS resolution. Under these conditions, we achieved near-baseline separation of KA and KAL using a range that encompassed only the analytes of interest (Figure 2A). Comparing mobilograms of pure standards confirmed the peak at 0.693 Vs/cm^2^ (147.7 Å²) to be KA and the peak at 0.713 Vs/cm^2^ (152.0 Å²) to be KAL.

It is worth noting that the use of a narrow mobility range relies heavily on accurate calibration of the 1/K_0_ range prior to data collection. Because it is challenging to find TIMS calibrants that span such narrow 1/K_0_ windows, we performed calibration over a broader range (0.60–1.30 Vs/cm²) using Agilent Tune Mix and subsequently reduced the mobility range for data collection. This method proved consistent, with minimal variation in the measured 1/K_0_ values for both KA and KAL across runs. To maintain consistency, we continued using a 1:1 KA and KAL mixture to tune TIMS parameters prior to data acquisition.

After optimizing the TIMS parameters via ESI, we translated our settings to a MALDI-TIMS-MS method for high-throughput screening. Following the evaluation of several matrices in negative ion mode, 1,5-diaminonaphthalene (1,5-DAN) was selected for MALDI analysis, as it was the only matrix that generated signal for *m/z* 212.09 (Figure S8). The TIMS parameters were transferred directly to the MALDI-TIMS-MS method, and tune parameters were subsequently adjusted to improve sensitivity for KA and KAL (Table S4). Notably, during DI-ESI of the 1:1 KA and KAL mixture, the signal intensity for KAL was significantly lower than that of KA (28% relative intensity), suggesting a difference in ionization efficiency between the two analytes under ESI conditions (Figure 2B). Interestingly, when the same 1:1 mixture was analyzed via MALDI-TIMS-MS, the intensities of KA and KAL were comparable (93% relative intensity for KAL), suggesting that MALDI ionization offers a more reliable platform for relative quantitation—particularly when KAL concentrations are low.

Despite narrowing the mobility range, considerable chemical noise persisted in our MALDI-TIMS-MS spectra—even with purified standards. We hypothesized that coupling quadrupole isolation with TIMS separation could further enhance signal and improve sensitivity for kainoids, particularly within complex biological matrices such as microbial colony material.^54^ Parallel reaction monitoring–parallel accumulation serial fragmentation (PRM-PASEF) has recently been described as a method that synchronizes quadrupole ion isolation with TIMS separation, enabling the targeted acquisition of multiple MS/MS spectra within a single TIMS ramp. Although primarily used for MS/MS, we postulated that PRM-PASEF could function as a highly targeted pseudo-MS^1^ method when operated at low collision energy to prevent fragmentation—thereby filtering out all ions except KA and KAL. The default collision cell voltage on the Bruker timsTOF fleX is 10 eV, even during MS¹ acquisition. By setting the PRM-PASEF collision energy table to 10 eV, the method effectively operates as an MS^1^ scan with both TIMS and quadrupole ion isolation. Under these conditions, MALDI-PRM-PASEF produced a substantial reduction in noise compared to standard MALDI-TIMS-MS (Figure 2C). We carried this improvement forward into our subsequent analyses of biological samples and mutagenic libraries.

### MALDI-TIMS-MSI for optimization of on-agar KabC expression

After optimizing a MALDI-TIMS-MS method to monitor KA and KAL, we turned to biological sample preparation. Mutagenic libraries are typically expressed in microbial hosts and assayed via whole-cell lysates or biotransformations, with scalable methods for KA production already reported.^26^ Although faster than purifying enzymes for in vitro assays, these approaches require multiple handling steps that can take several days, and the resulting mixtures usually require cleanup (e.g., chromatography or extraction) prior to high-throughput screening. Moreover, MALDI-MS is highly sensitive to ion suppression from salts and buffers in lysates, often necessitating additional cleanup prior to analysis, which we sought to address.^55^

Several methods have been developed for direct colony-based analysis of biotransformation reactions and secondary metabolite production across various MS modalities.^56–58^ Our lab contributed to one such platform: “IDBac”, a MALDI-TOF-MS-based tool designed to reduce taxonomic and chemical redundancy in the construction of microbial drug discovery libraries.^57^ Drawing on our experience with direct colony-transfer MS analysis,^59^ we adapted the sample preparation workflow from IDBac to enable monitoring of kainoid isomer formation directly from *E. coli* colonies spread onto MALDI target plates. We directly transformed *E. coli* expressing WT DsKabC and its Y347F variant (as a positive control for increased KAL production) onto LB agar plates containing 1 mM PKA substrate (Table S1), allowing protein expression and reaction conversion to proceed overnight.

Before adopting a colony-based approach, we used MALDI mass spectrometry imaging (MSI) to optimize on-plate induction and conversion, focusing on IPTG concentration and temperature. The T7 promoter in the pET28a(+) plasmid allows basal expression even without inducer, and although the lac operator largely mitigates this, some uninduced expression persists.^60^ To assess its impact, DsKabC was expressed both with 100 µM IPTG—based on established agar plate protocols—and without IPTG. Cultures were grown at 37 °C for 10 h to promote colony formation, then shifted to 30 °C for 6 h to support protein expression and product conversion. No conversion was observed at 18 °C, whereas incubation at 30 °C supported DsKabC expression and subsequent product formation. Interestingly, product ion intensity (*m/z* 212.09) was higher when IPTG was omitted, regardless of the temperature shift. The combination of IPTG omission and the temperature transfer produced the highest median intensity and greatest spread in values, suggesting enhanced product conversion and higher DsKabC expression, while the temperature shift alone increased variability irrespective of IPTG presence (Figures 3A-B, S9).

**Figure 3.**
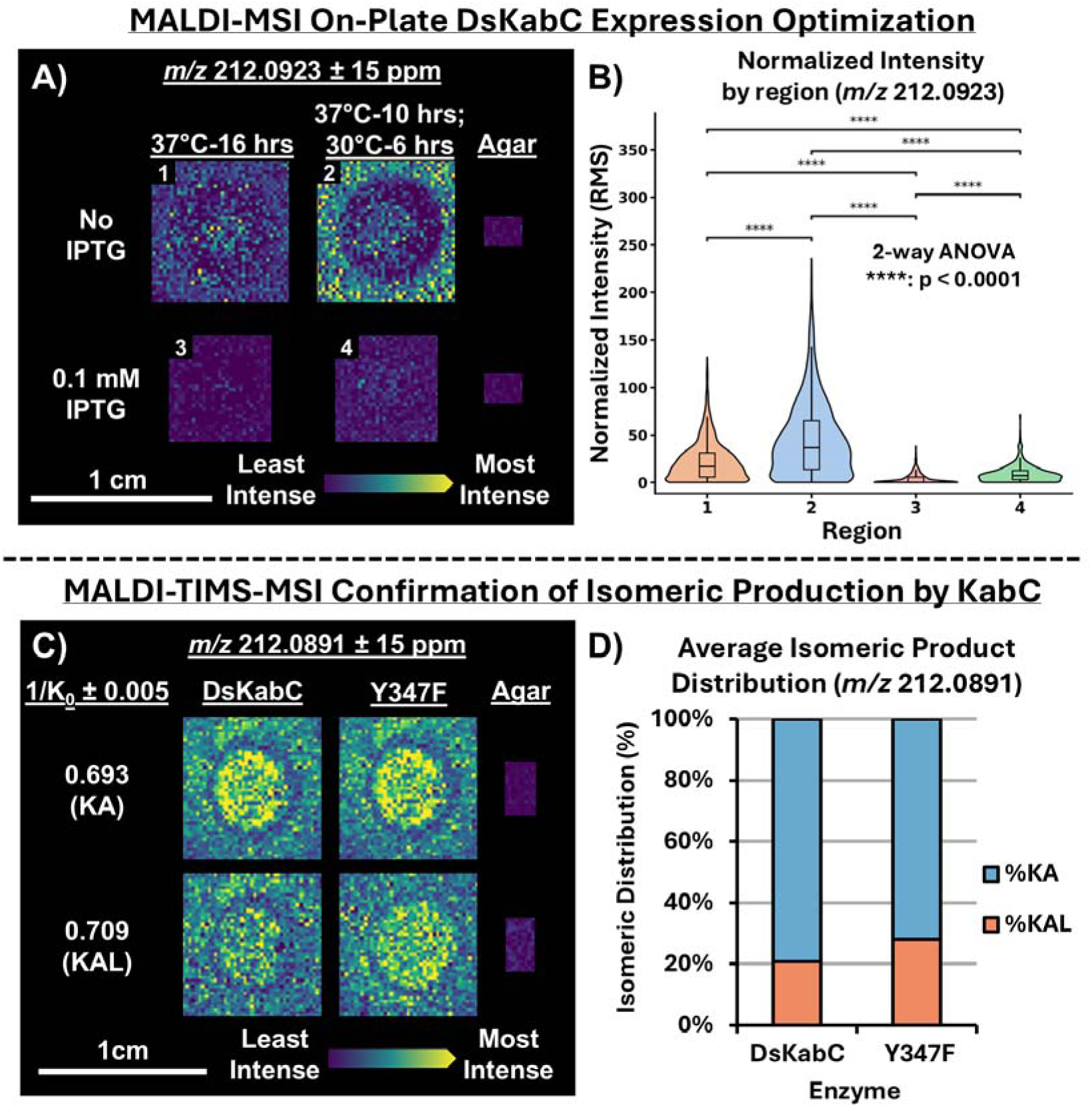
Optimization of on-agar DsKabC expression via MALDI-TIMS-MSI. A) Ion images of BL21 (DE3) *E. coli* expressing WT DsKabC, grown on LB/KAN agar containing 1 mM PKA substrate. The amount of IPTG and the temperature of incubation is varied between all four biological replicates and all images were collected during the same MALDI-MSI acquisition. B) Box and whisker plots highlighting the distribution of intensity values for *m/* 212.0923 normalized by root mean square (RMS). The box and whisker plots are overlaid with violin plots highlighting the density and spread of normalized intensity values within each ion image. Significance was determined using Tukey’s post-hoc test, following a two-way ANOVA with temperature and IPTG presence serving as the independent variables. C) Ion images of BL21 DE3 *E. coli* expressing either WT DsKabC or DsKabC Y347F, grown on LB/KAN agar containing 1mM PKA substrate (*m/z* 212.0891). D) The average relative abundance of KA and KAL across all individual spectra for both WT DsKabC (n = 3052) and DsKabC Y347F (n = 3744). Y347F continues to display increased production of KAL relative to WT DsKabC, consistent with the initial *in vitro* assessment of Y347F activity via LC-MS (Figure 1E).

MSI enabled a spatial, semi-quantitative analysis of kainoid isomers in *E. coli* expressing either wild-type DsKabC or the Y347F mutant. The mutant showed a modest but consistent increase in KAL relative to KA, consistent with LC-MS results (Figures 1E, 3C). In the TIMS range of 0.67–0.73 Vs/cm², two distinct signals at 0.693 (KA) and 0.709 Vs/cm² (KAL) matched prior standard analyses for *m/z* 212.09 (Figure 2). Ion images confirmed differences in KA and KAL signal intensities between WT and Y347F strains (Figure 3C). This trend was further supported by plotting the average relative abundance of KA and KAL across all spectra collected for both genotypes during the MALDI-TIMS-MSI experiment (Figure 3D). Ultimately, MALDI-(TIMS)-MSI proved effective for optimizing on-agar expression conditions and enabling robust relative quantitation of KA and KAL, establishing a foundation for a higher-throughput, colony-based MALDI-TIMS-MS screening platform.

With biological sample preparation optimized, the next challenge was adapting our workflow for high-throughput analysis. The key modification involved transferring biomass from agar plates onto MALDI target plates, rather than analyzing biomass directly on the agar surface. This approach reduces chemical background from the media, facilitates systematic cataloging of mutant colonies for organized library screening, and enables the rapid, high-throughput “dried droplet” format typical of MALDI-MS workflows—an advantage not available in single-sample MALDI imaging experiments.

### Generation of KabC chimeric library and high-throughput screening via MALDI-TIMS-MS

To build on the limited success of our rational mutagenesis approach, we next generated a random mutagenic KabC library using error-prone PCR (epPCR). Our initial epPCR library encompassed 736 KabC variants, but only modest improvements in relative KAL production were observed in first-pass screening (Figure S11–13). To drive a more substantial shift in kainoid isomer distribution, we turned to DNA shuffling, which has been successfully employed to improve activity and alter function in other Fe/αKG systems.^61,62^ The nucleotide sequence of *gfKabC* was optimized from 64.9% to 82.7% *dsKabC* identity to improve their compatibility at degenerate codons.^61^ The two sequences were subjected to DNA shuffling (Tables S5-S8) and used to generate a chimeric library of variants with shuffled *dsKabC* and *gfKabC* sequences (Figure 4A).

**Figure 4.**
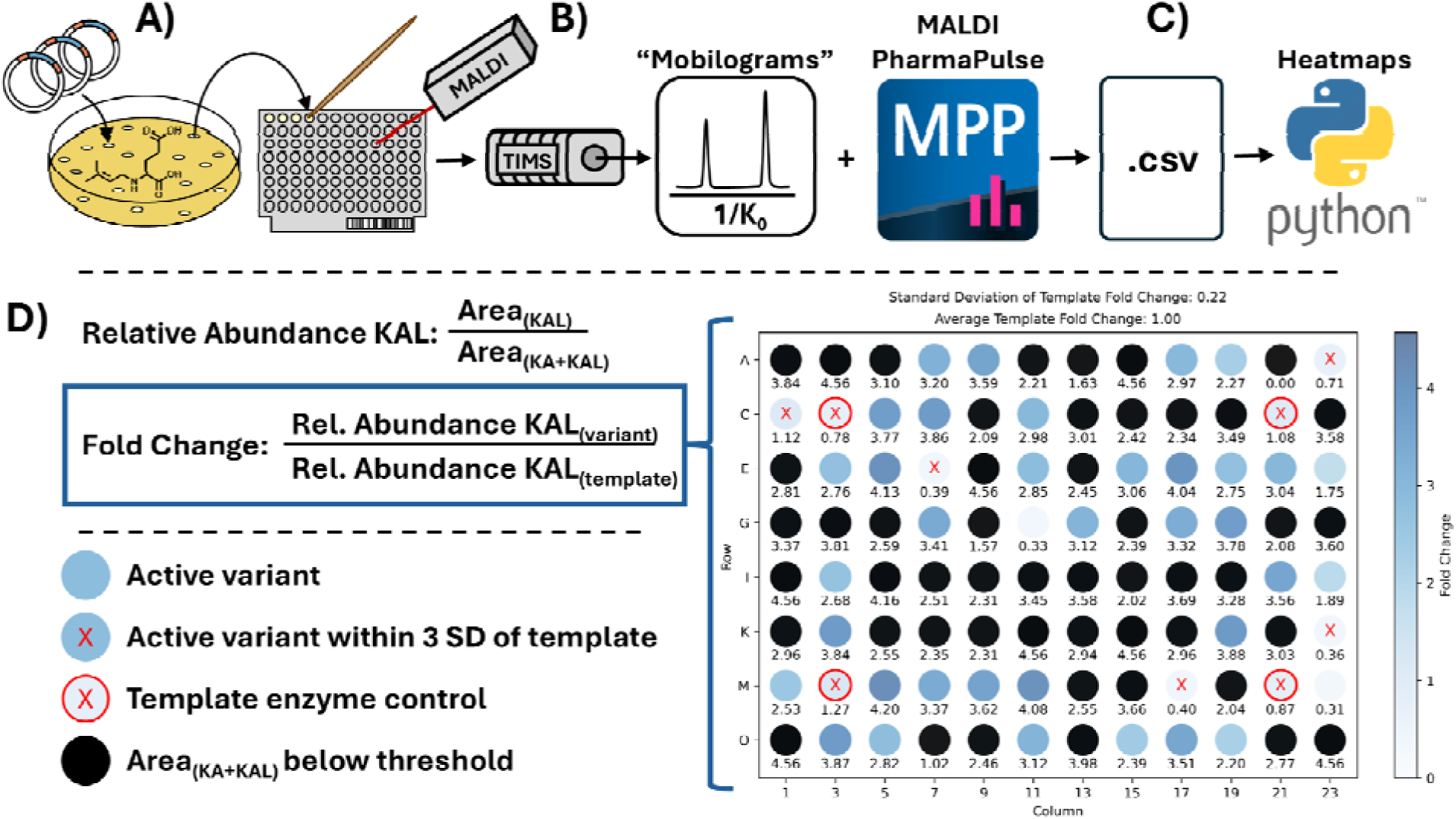
Schematic of MALDI-TIMS-MS screening platform. A) Following mutagenic PCR, the genes are ligated into a pET28a(+) expression plasmid and transformed into *E. coli*. The transformation plates contain 1 mM of PKA substrate to allow for KabC expression and substrate conversion in the same incubation period. The resulting colonies are directly transferred to MALDI plates, and matrix applied for downstream analysis. B) MALDI-TIMS-MS acquisition is performed, generating mobilograms that can subsequently be compiled into *.csv files using the MPP software. C) The generated *.csv files are fed into an in-house python script, generating heatmaps of variant fold change. D) The relative abundance of KAL for each variant and template controls is calculated first. The fold change is calculated by dividing the relative KAL abundance for each variant by the average of 4 DsKabC template control spots (red circle) for a 96-spot MALDI target. Blue indicates high fold-change, white represents zero-fold change, and black represents spots in which the total TIMS peak area for KA + KAL was less than a defined threshold (1000 was used for our screening). Variants within 3 standard deviations of the average relative KAL abundance of the template controls are marked with a red X.

To screen our chimeric KabC library in high throughput, we acquired MALDI-TIMS-MS data using the MALDI PharmaPulse (MPP) software, which is specifically designed for rapid evaluation of reactant and product screening campaigns via MALDI-MS (Figure 4B). To ensure robust analysis from single colonies, the MALDI laser was rastered randomly across each spot position (5x bursts) on the MALDI target to account for sample heterogeneity. The spectra from each raster point were summed into a consensus spectrum for each MALDI spot. The consensus spectrum was used for subsequent data analysis. The results were exported as a *.csv file, and an in-house Python script was used to visualize chimeric KabC activity as heatmaps of the fold change (FC) in relative KAL intensity with respect to the average of four DsKabC template enzyme control spots (DsKabC normalized to FC=1; Figure 4C-D).

In total we screened 318 variants (Figures S14, S15) from the shuffle library in <30 minutes (∼5 s per sample). DNA shuffling enriched for active variants, many of which had fold change values ≥4. For sequencing, we selected 6 variants with FC values >4, 10 variants with FC values between 3 and 4—potentially representing mutations that may lead to moderate changes in isomeric selectivity—and 2 variants with FC values <3 that, despite lower calculated fold changes, exhibited raw mobilograms suggestive of high relative KA production (Figure S16). When plotting the total intensity of all variants, 57 had total kainoid mobilogram peak areas greater than our selection threshold (Figure S15A). Subsequently plotting these 57 variants in terms of their relative KAL production revealed a clear split in the distribution of active variants, suggesting that 21 active variants primarily produced KA, 36 primarily produced KAL, and interestingly, none produced KA and KAL in near-equal measure (Figure S15B). All 18 variants selected for sequencing were broadly distributed across the 36 variants selective for KAL production. Of these 18 variants, 5 corresponded to wild-type *gfKabC,* while 7 were determined to have shuffled sequences. The remaining 6 hits yielded poor sequencing reads.

To confirm the activity of the shuffle library hits, each of the 7 chimeric enzymes was cloned into a pET28a(+) vector, heterologously expressed in *E. coli,* and purified via Ni-NTA affinity chromatography (Figure S17). Each purified variant was incubated *in vitro* with Fe^2+^, αKG, L-ascorbate, and PKA, and allowed to react for 18 hours in an aerobic environment.^24^ Following conventional LC-MS analysis (Figure S18, S19), five of the shuffled variants demonstrated improved KAL production (∼94 - >99% relative KAL abundance) compared to GfKabC (∼93% relative KAL abundance). Moreover, two variants (SL44 and SL306) exhibited a higher substrate consumption (>99% and ∼98%, respectively) compared to GfKabC (∼86%) (Figure 5A). The calculated relative abundance of KAL from the MALDI-TIMS-MS data correlated strongly with that calculated for the *in vitro* LC-MS data of the 7 tested shuffle variants (R^2^=0.95; Figure 5B). Notably, SL44 and SL306 were well-expressed with an N-terminal His tag, overcoming the solubility and expression limitations of GfKabC, which requires a maltose-binding protein (MBP) tag for adequate production in *E. coli*.^26^ SL306 retained activity at variable enzyme concentrations (0.5 - 5.0 mol%) compared to GfKabC and SL44 after storage at −70°C for 4 months, and multiple freeze-thaw cycles, suggesting increased stability over both GfKabC and SL44 (Figure S20).

**Figure 5.**
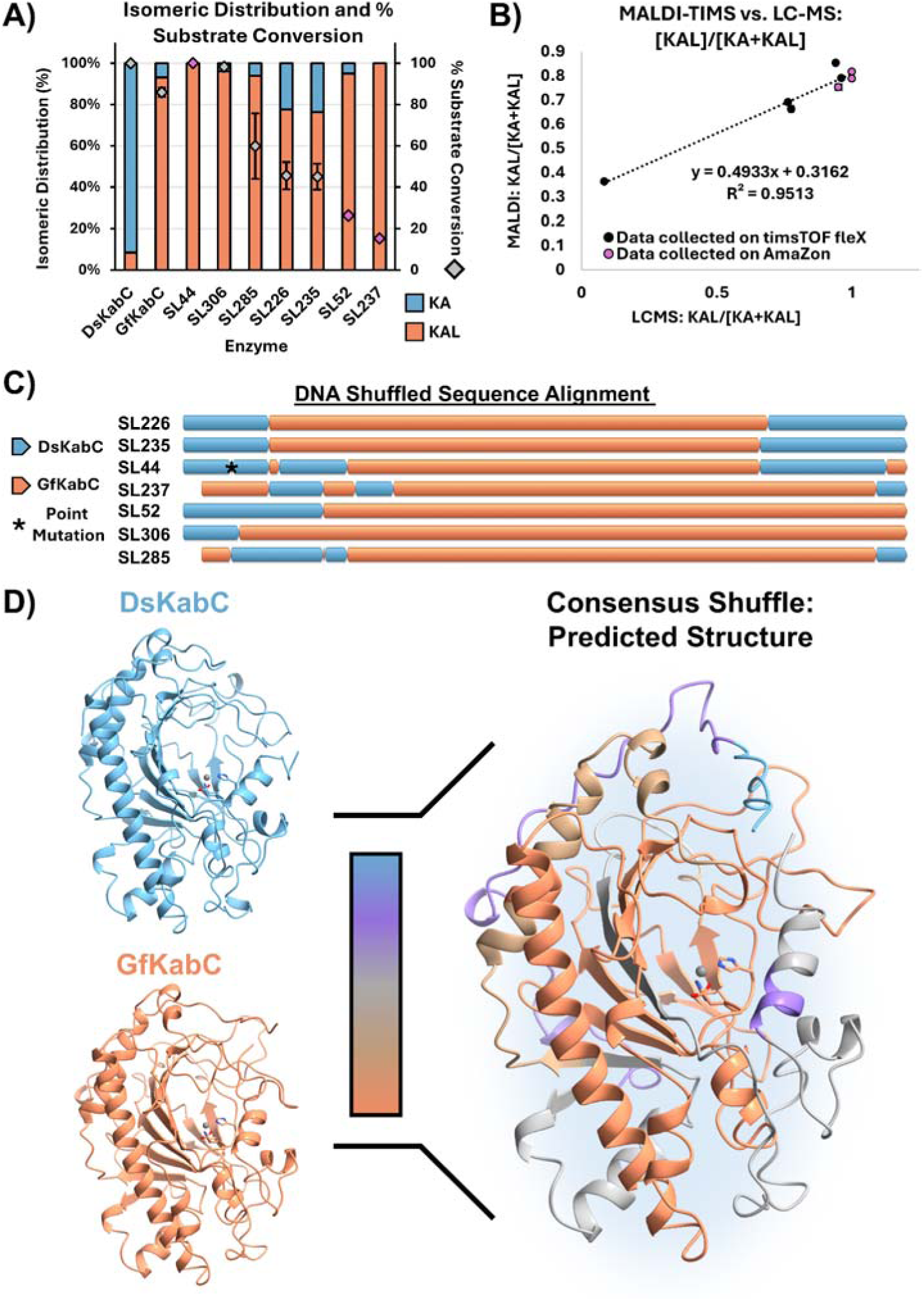
Overview of selected DNA-shuffled KabC variants. A) Isomeric distribution of KA and KAL represented as relative percent abundance, and percent substrate conversion for the selected DNA-shuffled variants determined via LC-MS *in vitro* assays. Grey diamonds represent data points collected using the timsTOF fleX mass spectrometer; pink diamonds represent datapoints collected using a Bruker AmaZon SL mass spectrometer. B) Correlation of measured relative KAL abundance between MALDI-TIMS-MS and LC-MS for the DNA shuffle variants studied *in vitro*. A high correlation was observed for KAL/[KA+KAL] between LC-MS and MALDI-TIMS-MS analysis on the seven purified shuffle library hits and WT DsKabC (R^2^ = 0.95). C) Simplified multiple sequence alignment schematic of the 7 shuffled variants. D) Comparison between the predicted structures of DsKabC (blue) and GfKabC (orange) with that of the consensus between all seven shuffled variants tested *in vitro*.

We generated AlphaFold 3^63^ models of all seven chimeric enzymes to investigate structural features that might explain differences in activity among the variants (Figures S21-22). Although no specific interactions were identified to account for the observed reaction outcomes, mapping regions of sequence conservation from DsKabC and GfKabC onto a predicted consensus AlphaFold 3 model allowed us to visualize conserved features across all seven shuffle variants (Figures 5C, 5D). The protein environment surrounding the active site was consistently derived from GfKabC in all seven shuffle variants that were high KAL producers. However, increased variability was observed at both the N- and C-termini, suggesting that the amino acid sequences in these regions are less responsible for product distribution, corroborating the rationally designed fusion constructs, *vide supra*. The enzymatic preference for KAL formation is likely influenced by complex secondary-sphere interactions that affect both the active-site geometry and the substrate binding pocket, as observed in other Fe/αKG enzymes.^61,64^ Efforts are underway to examine these interactions in greater detail to provide deeper mechanistic insight, identify structural determinants of KAL production, and leverage this for broader non-native kainoid biocatalysis.

## Conclusions

We have created and employed a high-throughput MALDI-TIMS-MS method for screening the activity of kainoid synthase biocatalysts on their native PKA substrate. Using this rapid assay that does not require enzyme purification, we generated and screened a 318-variant DNA shuffle library in ∼24 hours. This is a dramatic improvement over liquid-culture enzyme expression followed by LC-MS analysis, saving both time and labor. We identified seven KabC variants with improved conversion to KAL relative to GfKabC. Notably, two shuffled variants (SL44 and SL306) exhibited an almost complete shift toward KAL production, accompanied by enhanced substrate consumption, with SL306 exhibiting improved *in vitro* stability relative to GfKabC. Engineering kainoid synthases to mimic the isomeric output of GfKabC while retaining the favorable expression profile of DsKabC provides both a practical advantage for biocatalyst development and a robust model for analytical screening. Importantly, the combination of MALDI-TIMS-MS with enzyme engineering leverages the convenience and speed of direct colony MS analysis without sacrificing detailed analytical information, decreases reliance on LC, and enables the rapid assessment of large libraries for small molecule products. Further, PRM-PASEF enables the simultaneous analysis of multiple analytes with distinct mobility values, offering improved sensitivity and multiplexing capabilities, which is particularly advantageous for monitoring isomeric products. While demonstrated here for constitutional isomers, additional optimization can extend this platform to diastereomers and, with chemical derivatization, potentially enantiomeric products.^65^ Importantly, this strategy enables direct product readouts in enzymatic systems that are less compatible with conventional high-throughput screening (e.g., lacking chromophores, using non-chromogenic substrates, or requiring coupled assays). Finally, ongoing efforts to further engineer kainoid synthase variants will deepen our understanding of structure–function relationships and leverage this enzyme family for the construction of bioactive non-native kainoid natural products.

## Acknowledgements

We thank Arndt Asperger, Ph.D. and Heiko Neuweger, Ph.D. for helpful conversations regarding the MPP software. This work was supported by the National Institutes of Health (NIH) National Institute of General Medical Sciences of the NIH award R21GM148870 (LMS and SMKM) and training supplement (AJT), K12GM139185 (CAF), by National Science Foundation grants IOS-2220510 (LMS) and NSF graduate research fellowship program (RAS), and by the University of California Office of the President UCOP-UC LEADS program (MCJ).

## Data Availability

All MALDI and LC-MS data in this study are available under CC0 1.0 Universal License as raw (*.tdf) and open source (*.mzML) data formats. These data can be accessed here: doi:10.25345/C5SB3XB78; MassIVE accession: MSV000098944. The script used for this analysis along with documentation can be found at the following link: https://github.com/rashepherd/foldchangeviz.

## Supporting Information

The Supporting Information is available free of charge at https://pubs.acs.org/doi/XXX.

- Materials and experimental methods, media recipes, MSI sample preparation, all instrument tune and method parameters, chemical biology and biochemical procedures, protein purification data, LC-MS *in vitro* enzyme activity data, library heatmaps, and TIMS mobilograms for all variants highlighted in this manuscript.

