## Supplementary Information for "A High-Throughput Biocatalytic Platform for Screening Isomeric Kainoid Natural Products"

### Table of Contents

**Figure S1.** 13 sites targeted for site directed mutagenesis of DsKabC……………….............. S3
**Figure S2.** LC-MS chromatograms for whole cell lysate reactions of DsKabC SDM mutants.... S4
**Figure S3.** SDS-PAGE gels of batch purified DsKabC SDM mutants……………..…………… S5
**Figure S4.** LC-MS EICs for *in vitro* assays of batch purified DsKabC SDM mutants…………... S6
**Figure S5.** Alphafold models of KabC “fusion” enzyme constructs………………………………. S7
**Figure S6.** LC-MS EICs for *in vitro* assays of batch purified fusion enzyme constructs……….. S8
**Figure S7.** EIMs of purified fusion KabC *in vitro* activity assays………………………………….. S9
**Figure S8.** MALDI matrix evaluation in negative ion mode……………………………………….... S10
**Figure S9.** Replicate MALDI-MSI ion images of WT DsKabC and DsKabC Y347F……………. S11
**Figure S10.** Experimental reproducibility of KAL/[KA+KAL]..................................................... S12
**Figure S11.** Mutagenesis techniques used for this study……………………………….....….. S13
**Figure S12.** Heatmaps of variant fold change for KabC epPCR library………………………….. S14
**Figure S13.** epPCR Preliminary screening “hits”......................................................................... S15
**Figure S14.** Heatmaps of variant fold change for KabC DNA shuffle library…………………… S16
**Figure S15.** Assessment of total KabC DNA shuffle library activity via MALDI-TIMS-MS…….. S17
**Figure S16.** Colony-based EIMS of DsKabC and all seven purified DNA shuffle variants…… S18
**Figure S17.** SDS-PAGE gels of *E. coli* BL21(DE3) expressing shuffle library variants……..… S19-S24
**Figure S18.** *In vitro* EICs of DsKabC, GfKabC, and all seven purified shuffle variants……….. S25
**Figure S19.** LC-MS conversion data for WT KabC and shuffle variants……………………….. S26
**Figure S20.** *In vitro* stability assessment of GfKabC, SL44, and SL306……………………….. S27
**Figure S21.** Multiple sequence alignment of 12 sequenced hits…………………………………. S28-S29
**Figure S22.** Alphafold 3 models of the seven purified KabC shuffle library variants…………… S30
**Table S1.** LB+KAN+IPTG+PKA media recipe…………………………………………………….. S31
**Table S2.** Software information…...……………………………………………………………….. S32
**Table S3.** MALDI PharmaPulse Campaign Parameters ……………………………………...... S33
**Table S4.** Instrument Parameters for MALDI-TIMS-MS…………………………………………… S34-S36
**Table S5.** Primers used in this study……………………………………………………………… S52-S53
**Table S6.** Components of the PCR performed to obtain linear DNA for shuffling……………….. S54
**Table S7.** Components of the PCR performed to reamplify the reassembled DNA fragments.. S55
**Table S8.** Components of the PCR performed to obtain linearized pET28a(+) DNA………..… S56

**Experimental Methods and Procedures**……………………………………………………….. S37-S42
**Molecular Biology and Biochemical Methods**………………………………………………… S43-S56
**Codon-optimized DNA and Protein Sequences**………………………………………………. S44-S45

**13 divergent residues**

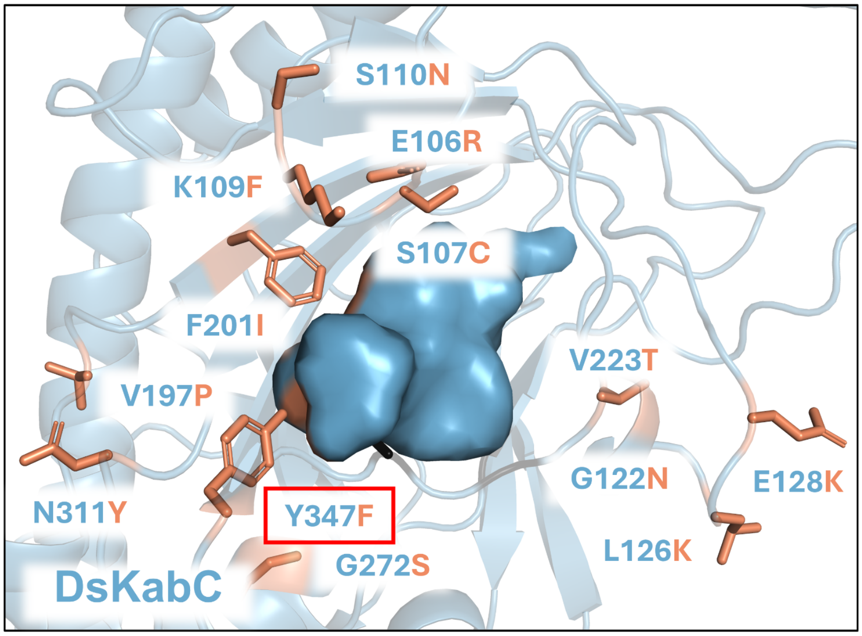


| Ds # | DsKabC | GfKabC | ReKabC | PpKabC | PhKabC |
|---|---|---|---|---|---|
| 106 | E | R | H | H | H |
| 107 | S | C | H | H | H |
| 109 | K | F | V | V | V |
| 110 | S | N | N | N | N |
| 122 | G | N | G | G | G |
| 126 | L | K | M | K | K |
| 128 | E | K | K | K | K |
| 197 | V | P | S | S | S |
| 201 | F | I | F | F | F |
| 223 | V | T | V | V | V |
| 272 | G | S | A | A | A |
| 311 | N | Y | H | H | H |
| 347 | Y | F | Y | Y | Y |

**Figure S1.** 13 divergent sites targeted for site directed mutagenesis determined using multiple sequence alignment. All targeted residues either diverge between Ds- and GfKabC, the broader ortholog set, or both. Only seven were successfully generated and screened for activity via lysate reactions (Figure S2). Highlighted in red is the SDM that was used as an additional positive control in screening development.

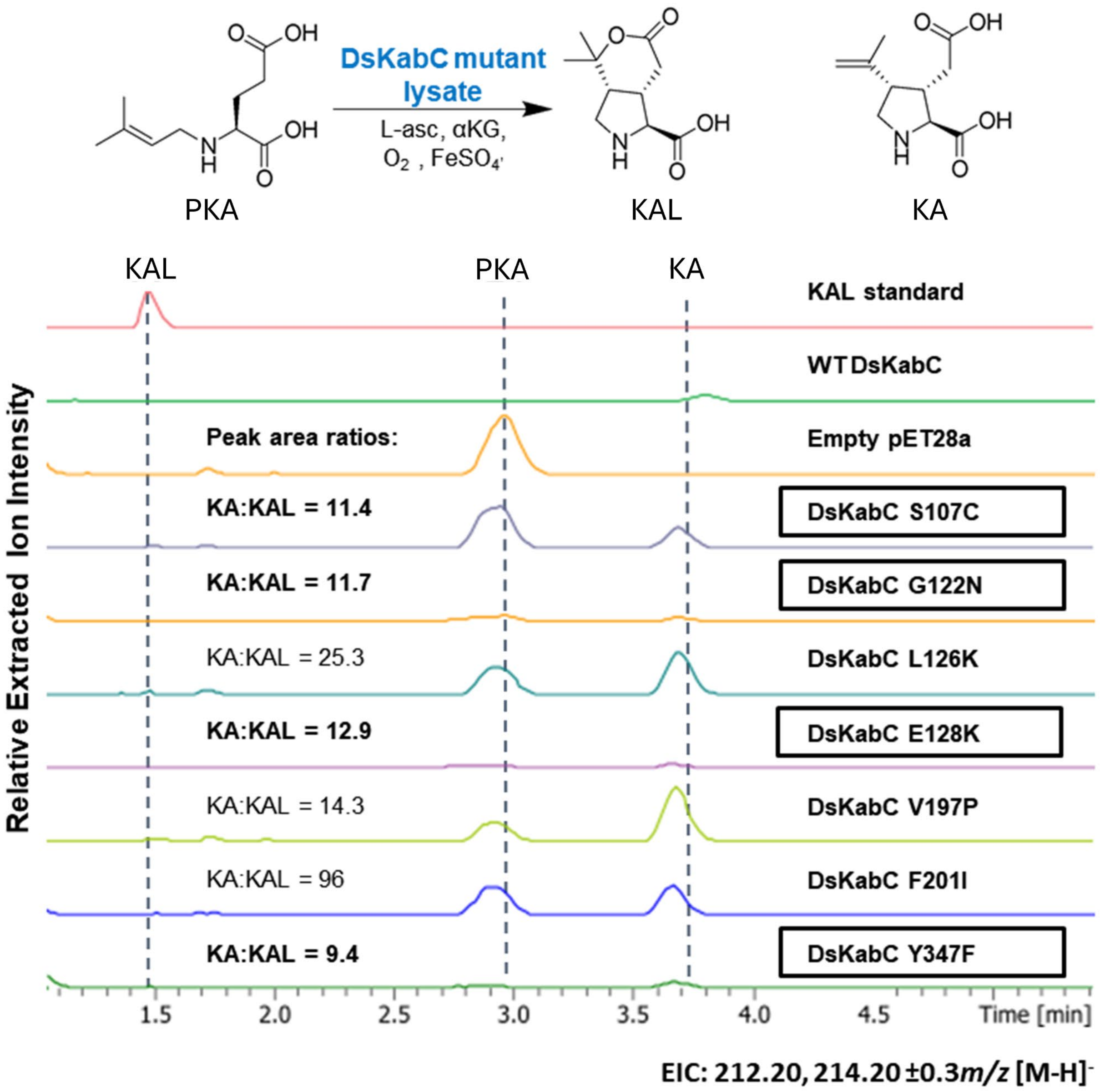


**Figure S2.** LC-MS extracted ion chromatograms (EIC) of whole-cell lysate reactions of the 7 successfully generated point mutants of DsKabC. 4 mutants were selected based on their higher relative abundance of KAL (as denoted by a lower number for the KA:KAL ratio and boxed in black).

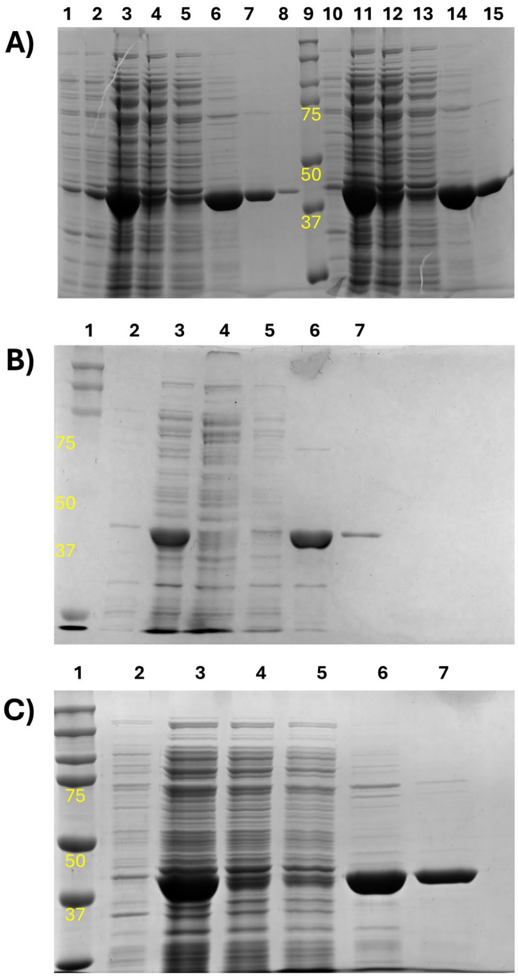


**Figure S3.** SDS-PAGE gels (Tris-glycine) of *E. coli* BL21(DE3) expressing DsKabC SDM mutants following batch purification. **Expected size of G122N: 44.0 kDa**. **A)** DsKabC G122N: Lanes (1-2) pellet (3) clarified lysate, (4) flow-through, (5) wash 100% A, (6) wash 50% B, (7-8) wash 100% B, (9) Precision Plus Protein Dual Color Standards ladder (Bio-Rad); DsKabC E128K: Lanes (10) pellet, (11) clarified lysate, (12) flow-through, (13) wash 100% A, (14) wash 50% B, (15) wash 100%. **B)** DsKabC Y347F: Lanes (1) Precision Plus Protein Dual Color Standards ladder (Bio-Rad), (2) pellet, (3) clarified lysate, (4) flow-through, (5) wash 100% A, (6) wash 50% B, (7) wash 100% B. **C)** DsKabC S107C: Lanes (1) Precision Plus Protein Dual Color Standards ladder (Bio-Rad), (2) pellet, (3) clarified lysate, (4) flow-through, (5) wash 100% A, (6) wash 50% B, (7) wash 100% B.

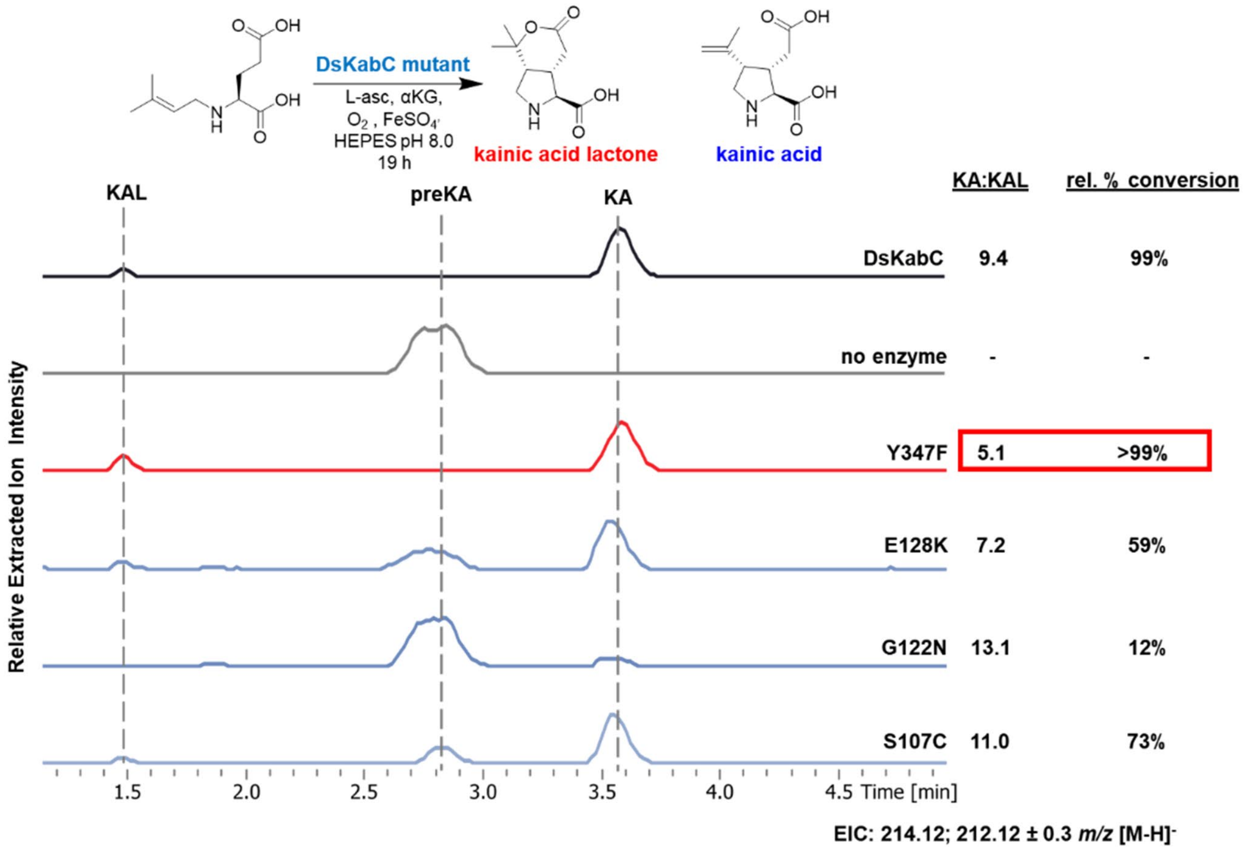


**Figure S4.** LC-MS EICs of purified *in vitro* enzyme reactions of the 4 prioritized single-point mutants of DsKabC. Y347F exhibited the highest relative abundance and relative percent conversion out of any other prioritized single-point mutant, boxed in red.

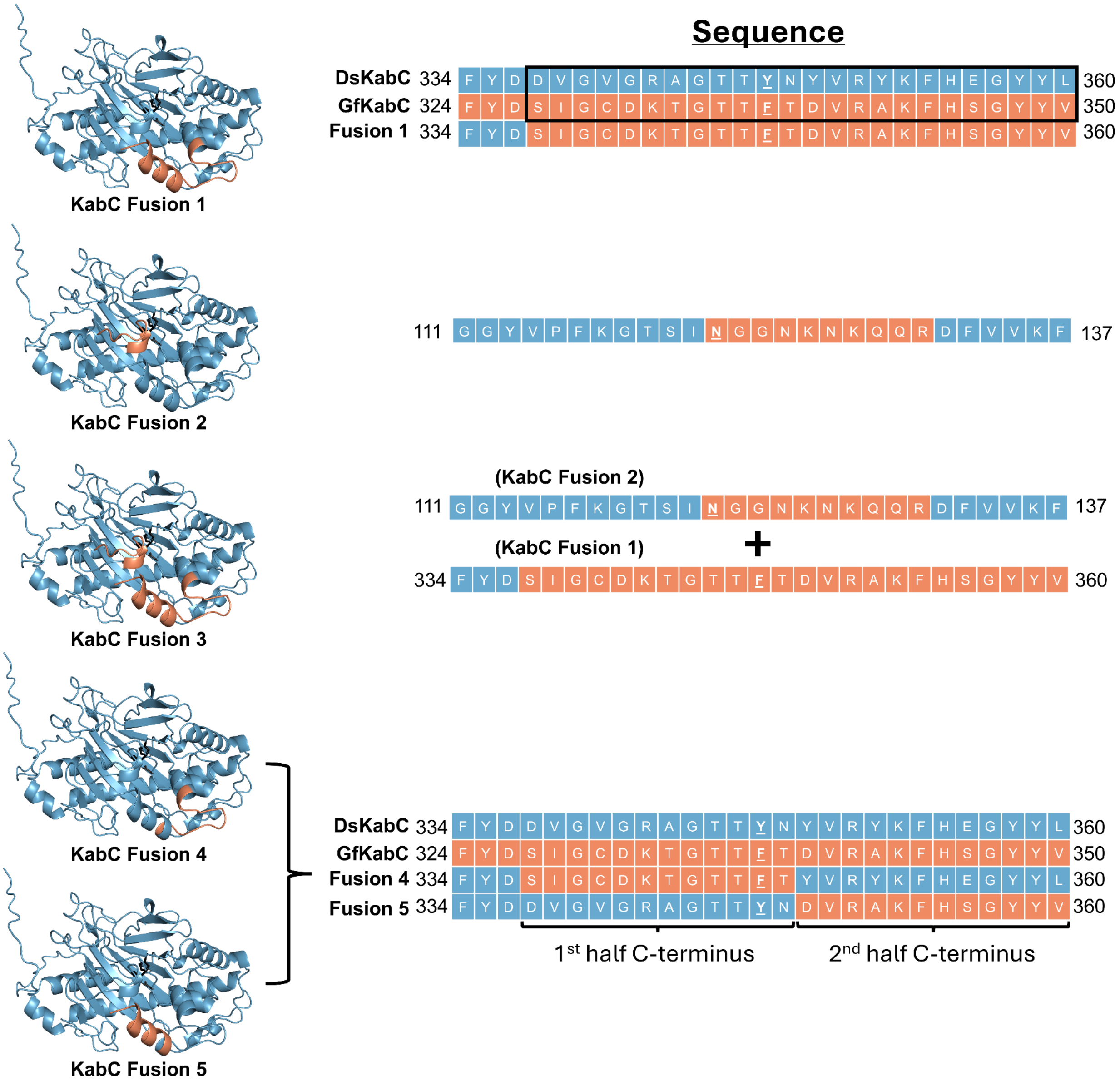


**Figure S5.** Alphafold models of the structures and relevant sequence portions of the 5 generated fusion enzyme constructs between Ds-(blue) and GfKabC (orange).

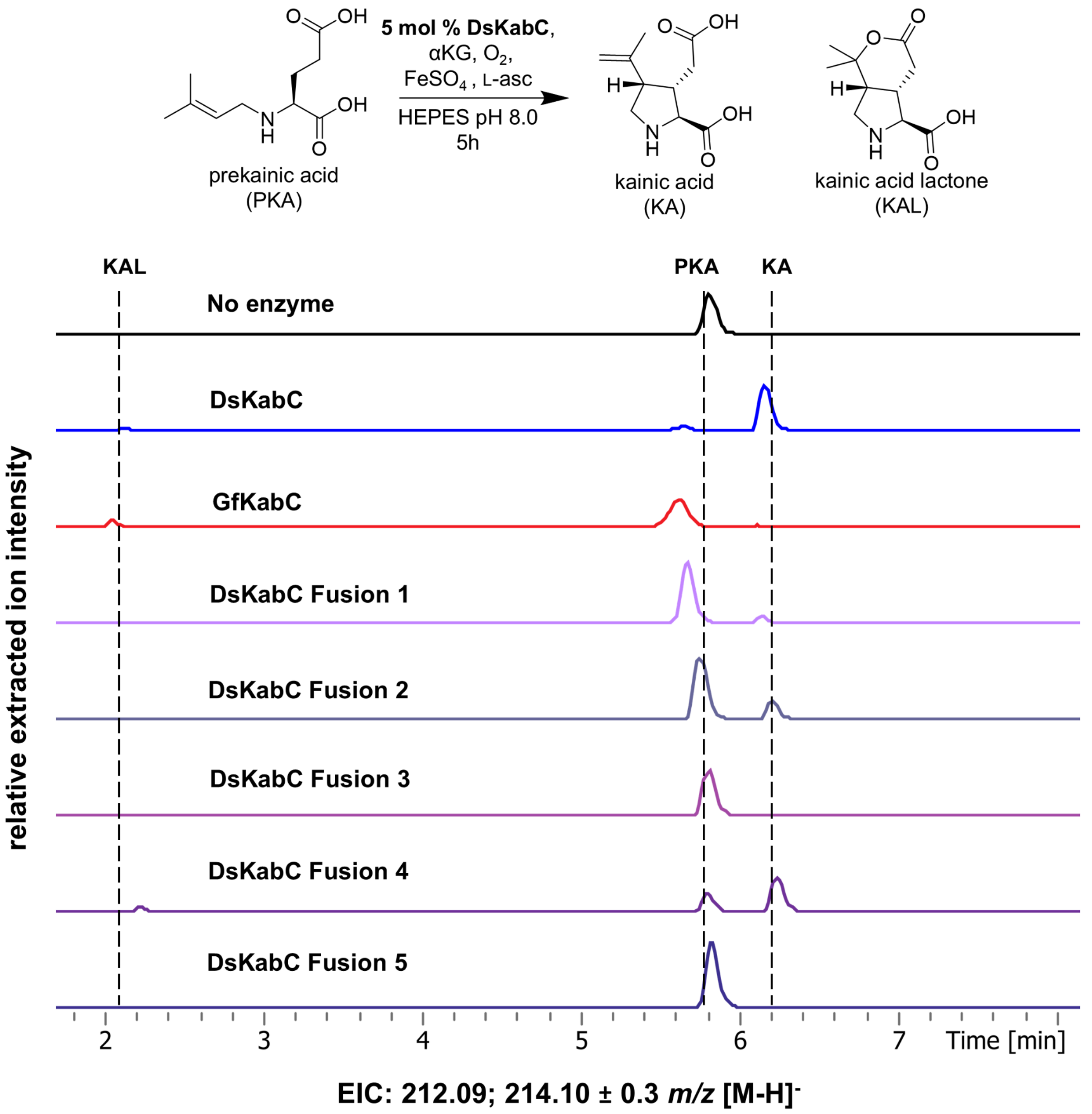


**Figure S6.** LC-MS EICs of purified fusion KabC enzyme *in vitro* activity assays. Only fusion 4 was observed to have a modest increase in KAL production, but substrate conversion was less than that of WT DsKabC.

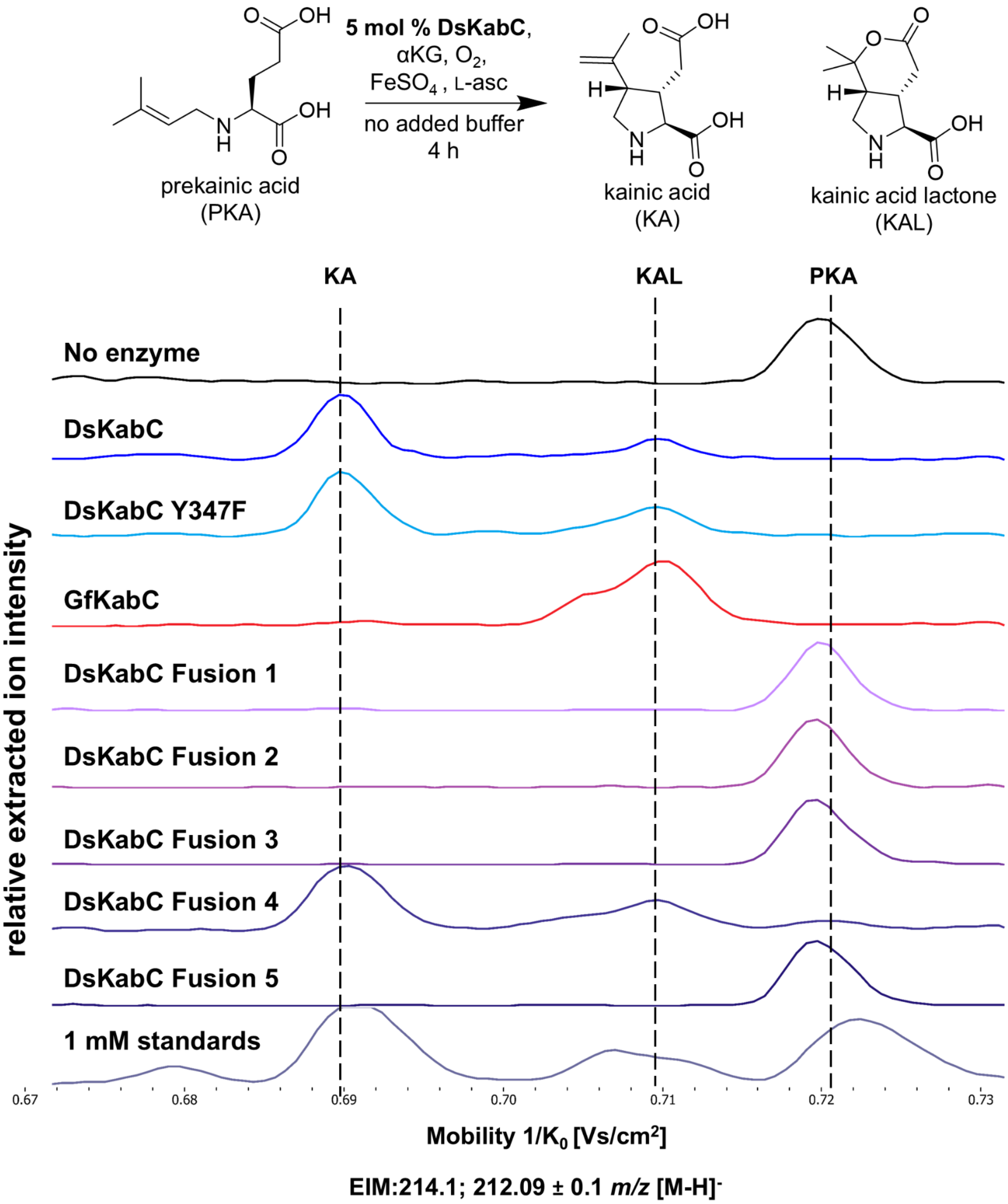


**Figure S7.** MALDI-TIMS-MS extracted ion mobilograms (EIMs) of purified fusion KabC *in vitro* activity assays (performed in water instead of HEPES buffer to prevent ion suppression from salts). Only fusion 4 was observed to have a modest increase in KAL production, consistent with the corresponding LC-MS data.

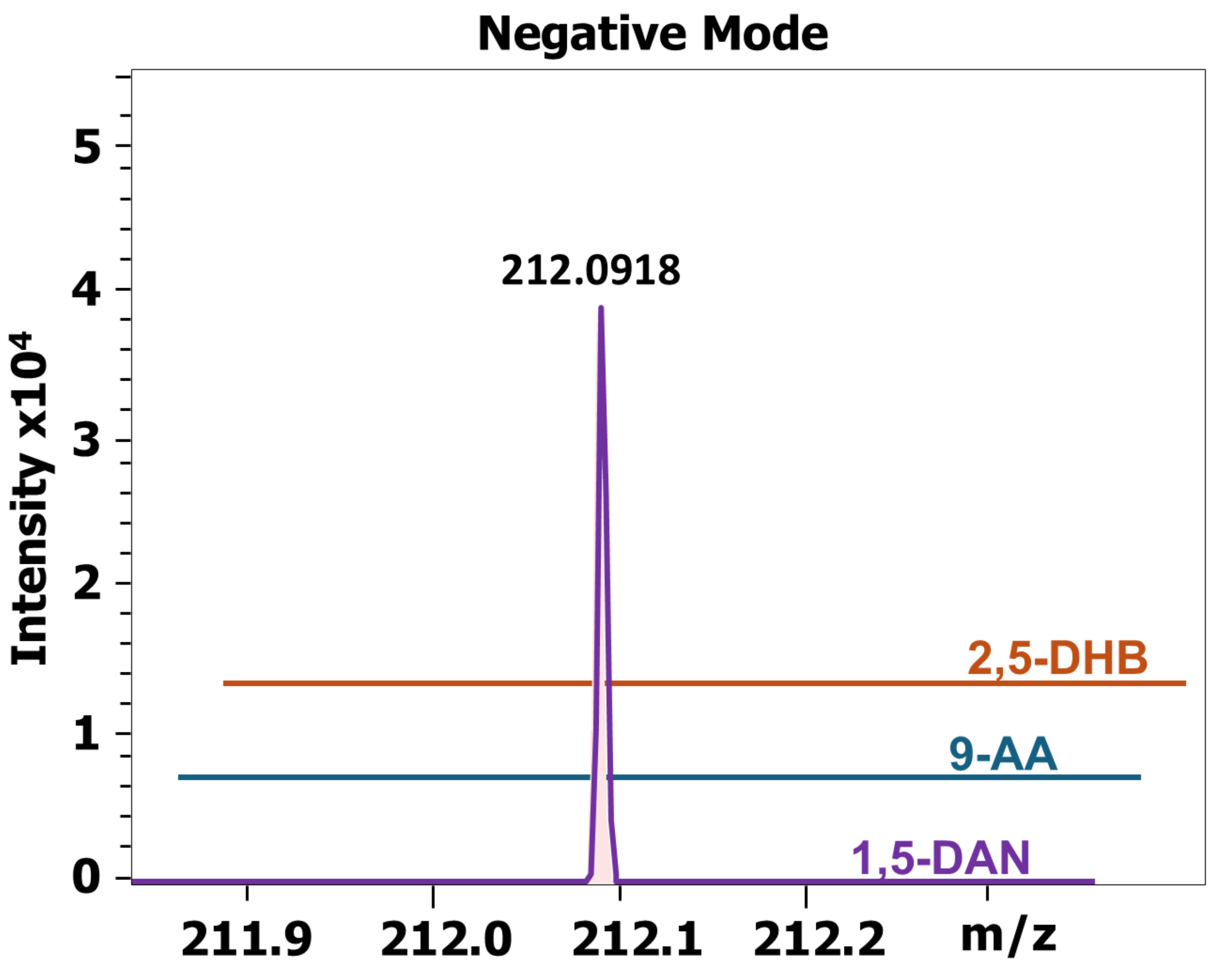


**Figure S8.** MALDI matrix evaluation in negative mode for 1 mM KA + KAL with 1,5-diaminonaphthalene (1,5-DAN), 9-aminoacridine (9-AA), and 2,5-dihydrobenzoic acid (2,5-DHB). 1,5-DAN is the only matrix that provided signal for KA and KAL.

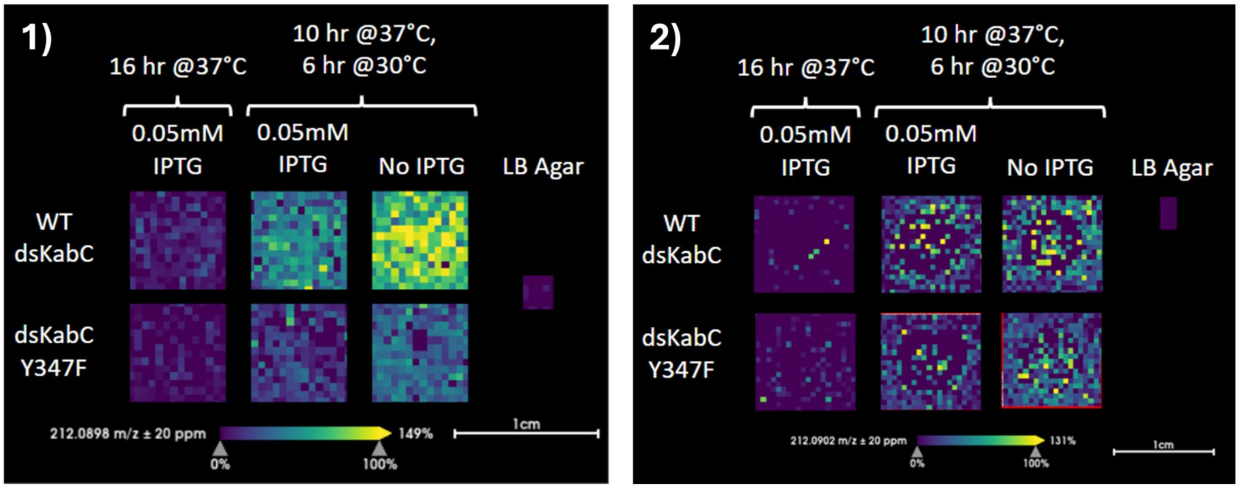


**Figure S9.** Replicate MALDI-MSI ion images of BL21 (DE3) *E. coli* expressing WT DsKabC at different incubation temperature conditions and different IPTG concentrations. Note that TIMS was not used for these analyses, as production of both KA and KAL were being monitored.

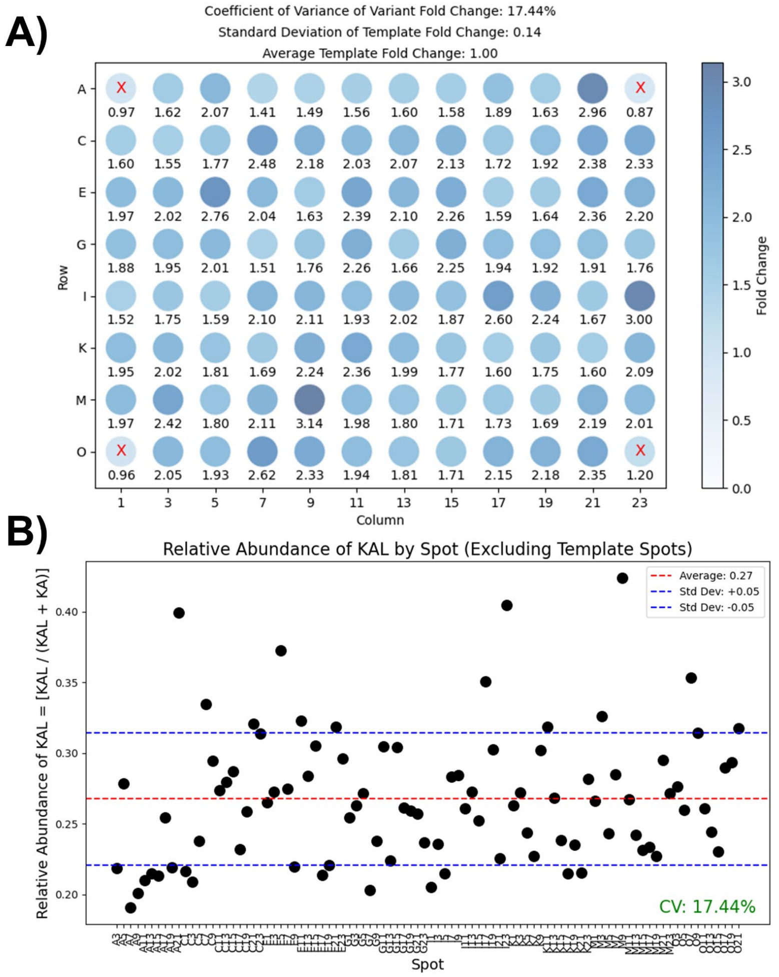


**Figure S10.** Experimental reproducibility of KAL/[KA+KAL] across 92 dsKabC Y347F colony replicates used in MALDI-TIMS-MS analysis. A) Heatmap displaying KAL/[KA+KAL] for DsKabC Y347F relative to that of WT DsKabC (control spots are indicated by red X's and were selected by the timsMAP Python script QC threshold). B) The distribution of KAL/[KA+KAL] across the 92 DsKabC Y347F replicates displayed in panel A. The coefficient of variation (CV) was determined to be 17.44%, which is sufficient for variant prioritization, and indicates MALDI-TIMS-MS can adequately distinguish between colonies with differing kainoid production. Notably, none of the DsKabC Y347F spots were within 3 standard deviations of the average activity of the template DsKabC controls (controls highlighted by the red X's).

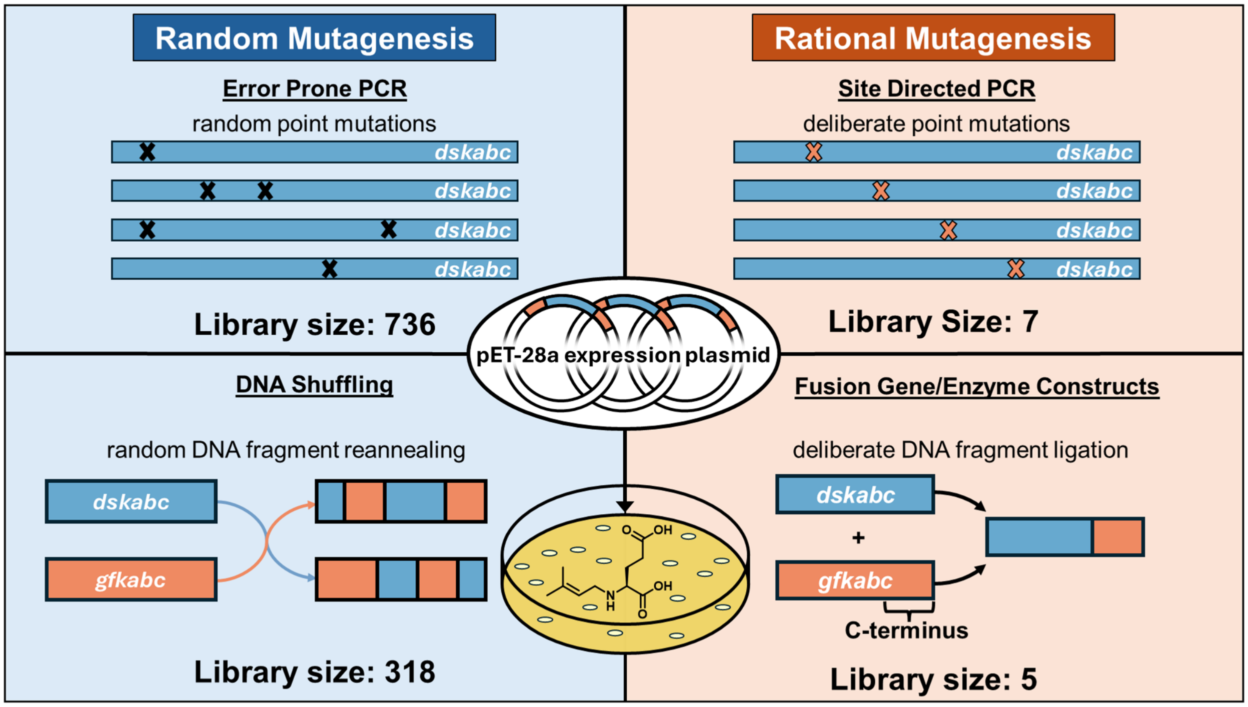


**Figure S11.** Depiction of the various mutagenesis techniques used in this study and their respective library sizes. In total, 1,054 random KabC variants were screened using the MALDI-TIMS-MS screening platform, representing ~ 1.5 hours of total instrument acquisition time.

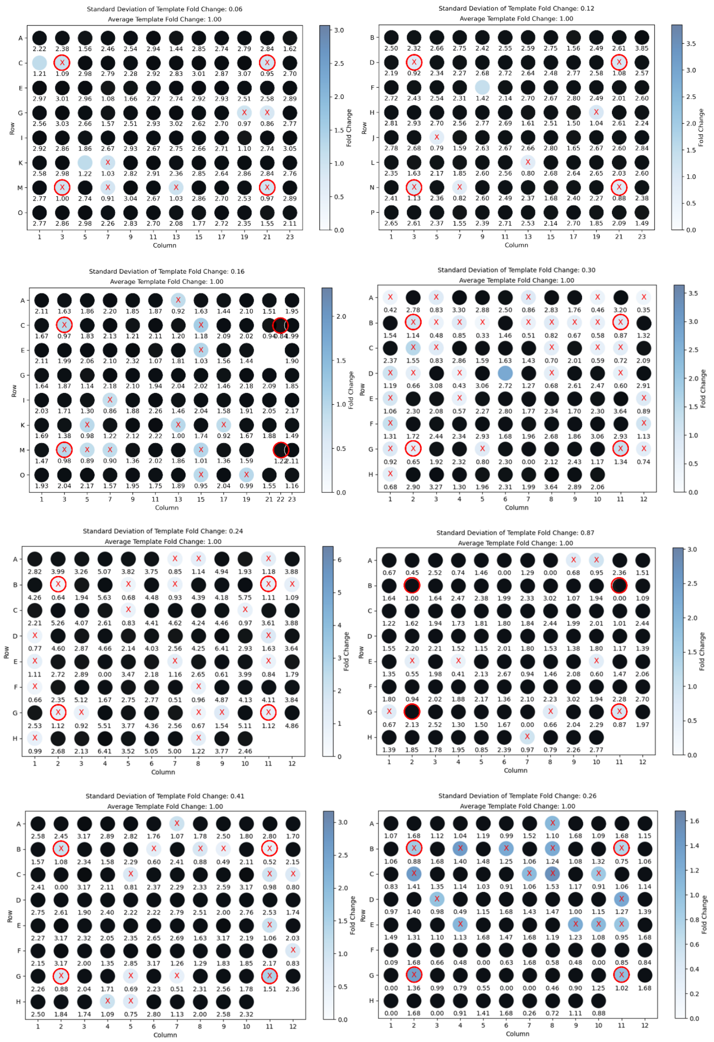


**Figure S12.** Heatmaps of variant fold change for KabC error prone mutagenesis library. Maximum fold change is denoted as blue, and a fold change of zero is denoted as white. Black denotes spots where the total TIMS peak area is below the set threshold (1000 during our screening campaign) to reduce the selection of false positives (variants that do not convert to KA and KAL have no product signal to measure and are therefore unreliable). Red X's represent spots in which the fold change in relative KAL abundance is within 3 standard deviations of the average of 4 DsKabC Y347F control spots (red circles).

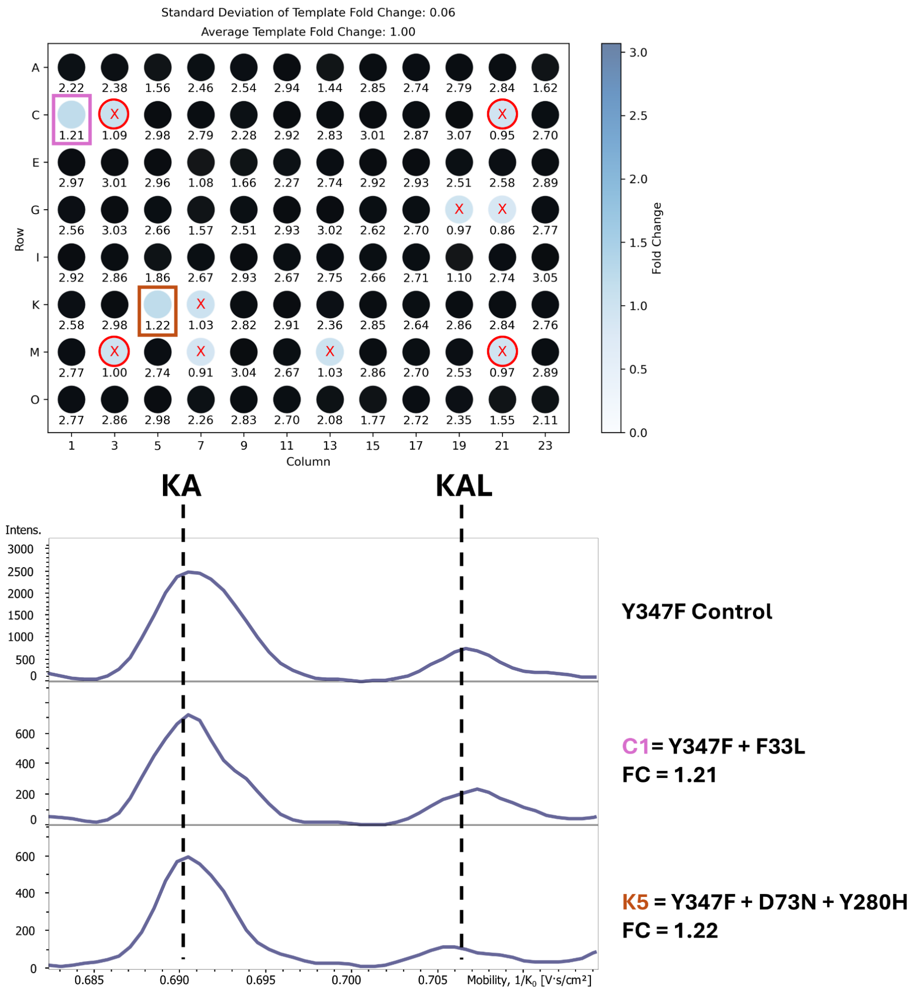


**Figure S13.** Two ‘hits’ from KabC epPCR library screening with corresponding mobilogram data. Two hits were chosen for DNA sequencing following MALDI-TIMS-MS screening of the generated KabC epPCR library. The variant at C1 was determined to have an additional point mutation at F33L, while the variant at K5 underwent two additional point mutations (D73N and Y280H). Since only modest changes in relative KAL abundance were observed for variant spots C1 and K5 (FC=1.21 and 1.22, respectively) neither were carried forward for *in vitro* validation via LC-MS.

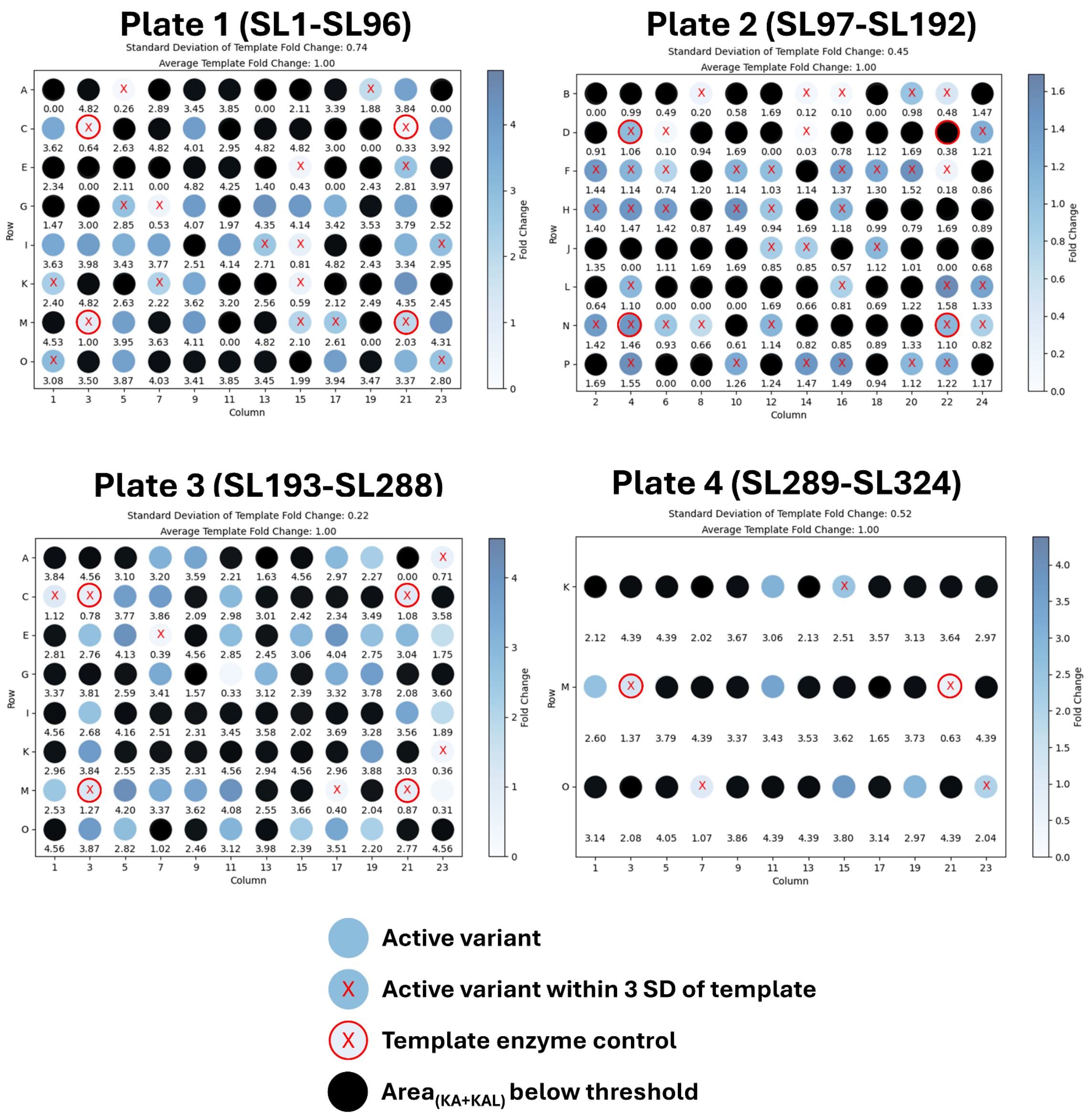


**Figure S14.** Heatmaps of variant fold change for the DNA shuffle library. Maximum fold change is denoted as blue, and a fold change of zero is denoted as white. Black denotes spots where the total TIMS peak area is below the set threshold (1000 during our screening campaign) to reduce the selection of false positives (variants that do not convert to KA and KAL have no product signal to measure and are therefore unreliable). Red X's represent spots in which the fold change in relative KAL abundance is within 3 standard deviations of the average of 4 DsKabC control spots.

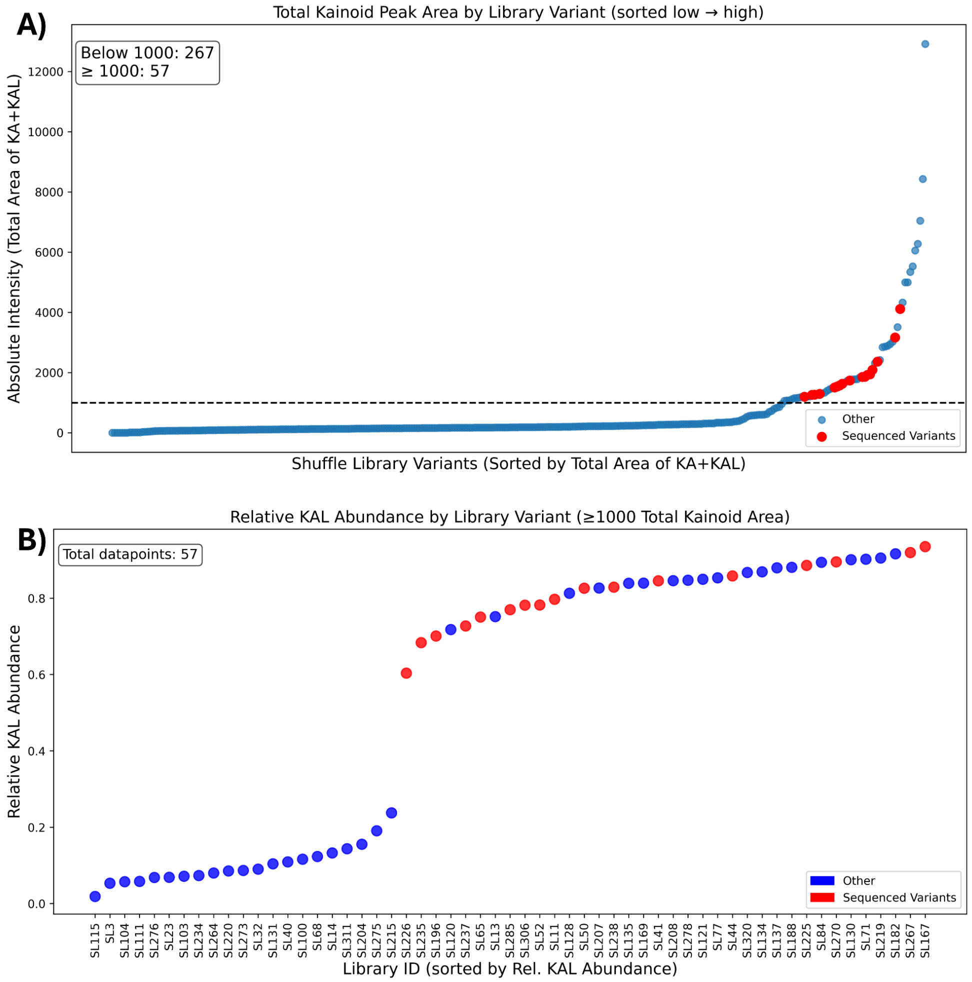


**Figure S15.** Assessment of total KabC DNA shuffle library activity via MALDI-TIMS-MS analysis. A) Scatterplot depicting all shuffle library variants ranked from lowest to highest absolute kainoid mobilogram peak area (extracted for *m/z* 212.09). The black dotted line represents the total peak area threshold for selection of active variants and/or variants with adequate signal-to-noise (1000 in this experiment). B) Scatterplot depicting the 57 shuffle library variants above the peak area threshold in panel A  ranked from lowest to highest relative KAL abundance (KAL/[KA+KAL]). In both plots, red indicates the 18 variants ultimately selected for DNA sequencing, and blue represents all other variants and control spots.

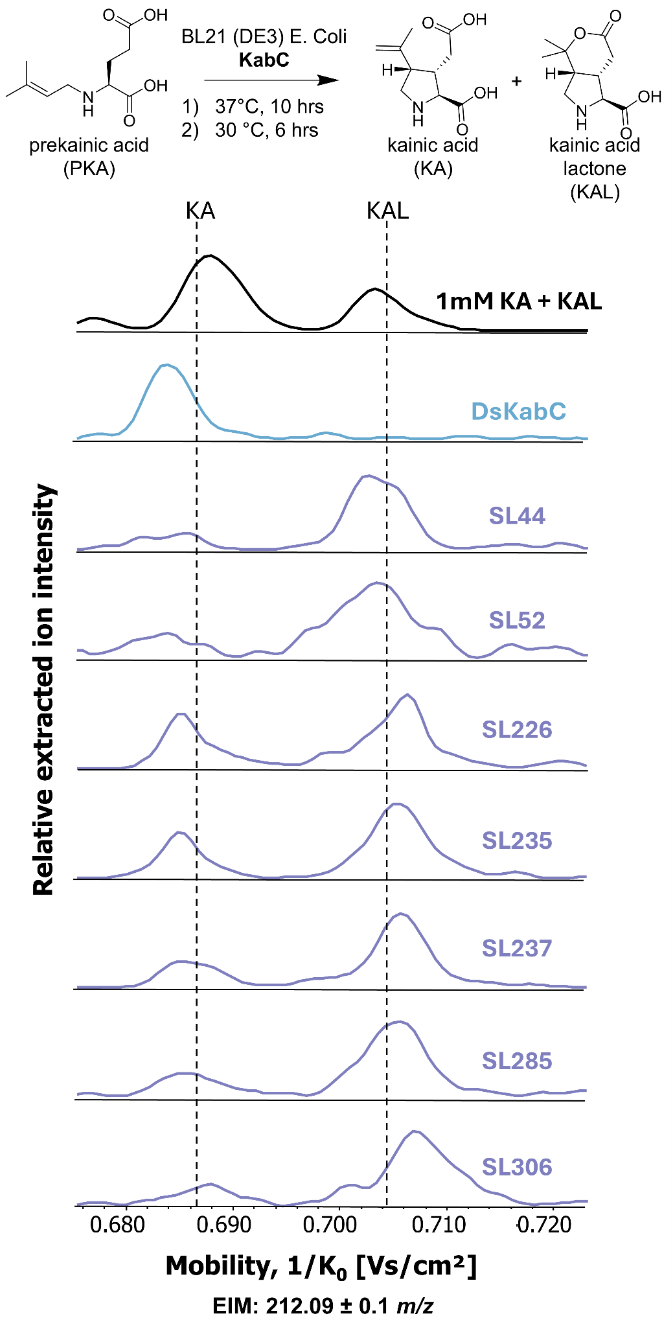


**Figure S16.** Colony-based EIMs of DsKabC and all seven DNA shuffled variants selected for subsequent *in vitro* characterization.

A) DsKabC purification

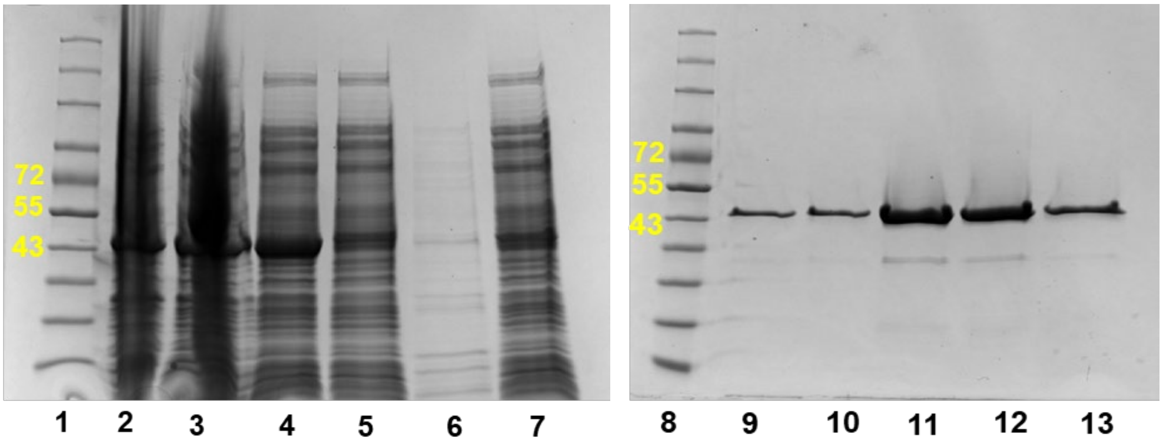


B) GfKabC purification

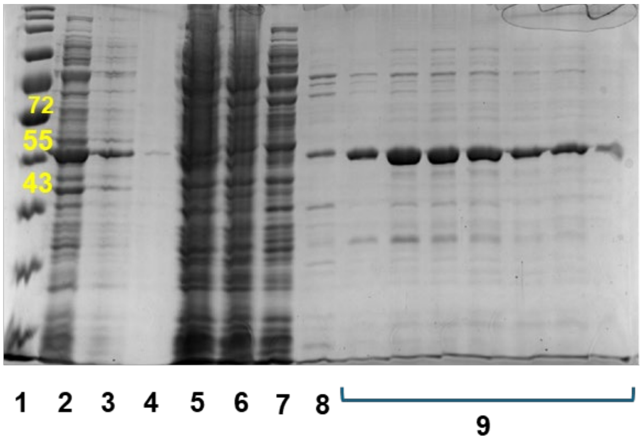

C) SL44 purification

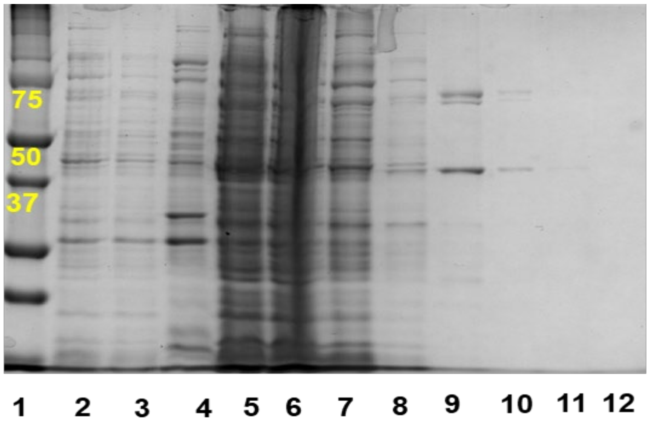


D) SL52 purification

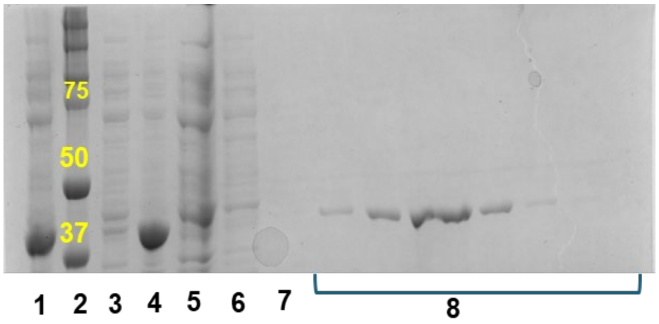


E) SL226 purification

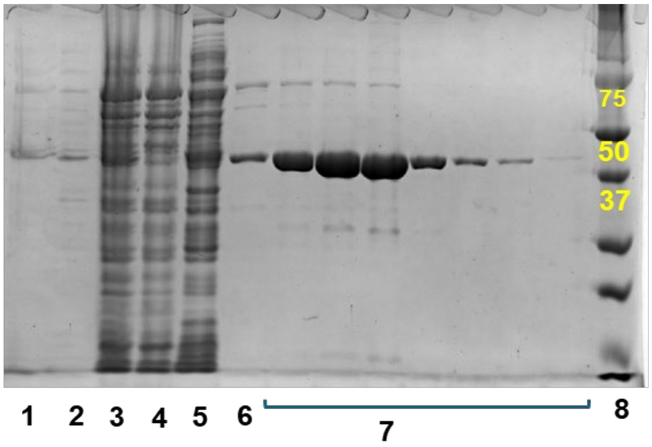

**F) SL235 purification**

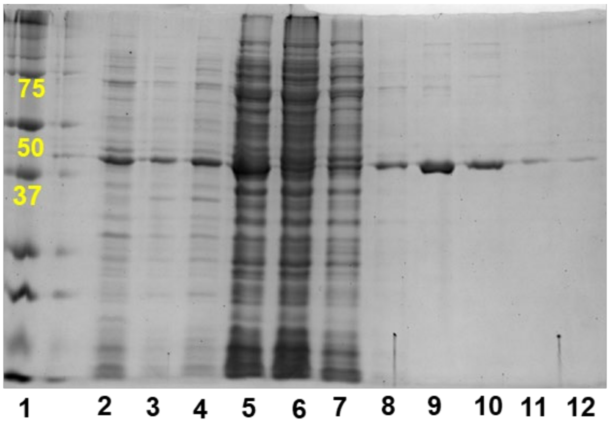


**G) SL237 purification**

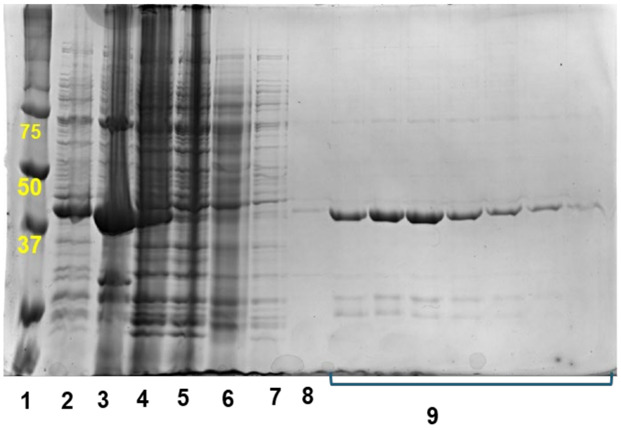

#### H) SL285 purification

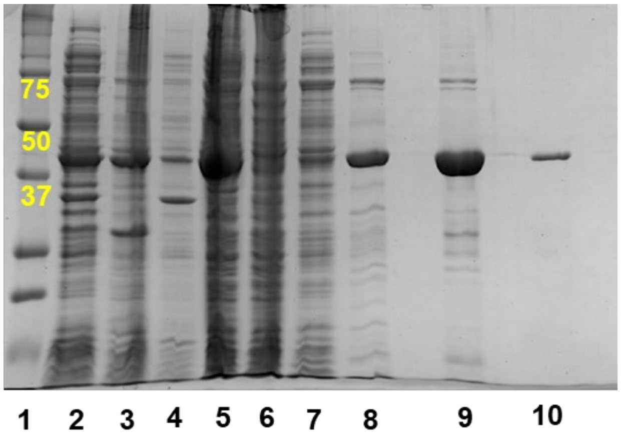


#### I) SL306 purification

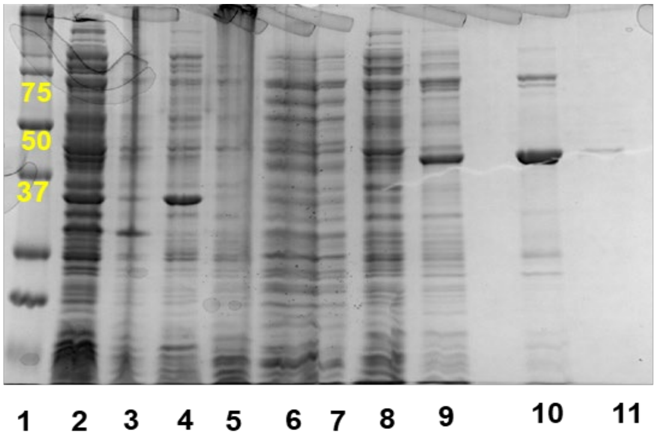


**Figure S17. A) DsKabC**: SDS-PAGE gels (Tris-glycine) of *E. coli* BL21(DE3) expressing DsKabC following FPLC purification. **Expected size of DsKabC: 43.3 kDa**. Lanes (1) and (8) Color Prestained Protein Standard Ladder (NEB), (2) pellet, (3) crude lysate, (4) clarified lysate, (5) fraction 4, (6) fraction 5, (7) fraction 8, (9) fraction 20, (10) fraction 22, (11) fraction 24, (12) fraction 26, (13) fraction 28. **B) GfKabC**: SDS-PAGE gels (Tris-glycine) of *E. coli* BL21(DE3) expressing GfKabC following FPLC purification. **Expected size of GfKabC: 42.4 kDa**. Lanes (1) Color Prestained Protein Standard Ladder (NEB), (2) pre-solubilized pellet, (3) crude lysate, (4) pellet,

(5) clarified lysate, (6) flow-through, (7) wash A, (8) wash 10% B, (9) fractions 4, 6, 8, 10, 12, 14, 17. **C) SL44**: SDS-PAGE gels (Tris-glycine) of *E. coli* BL21(DE3) expressing SL44 following batch purification. **Expected size of SL44: 41.5 kDa**. Lanes (1) Precision Plus Protein Dual Color Standards ladder (Bio-Rad), (2) pre-solubilized pellet, (3) crude lysate, (4) pellet, (5) clarified lysate, (6) flow-through, (7) wash A, (8) wash 10% B, (9) wash 50% B-1, (10) wash 50% B-2, (11) wash 100% B-1, (12) wash 100% B-2; **(D) SL52:** SDS-PAGE gels (Tris-glycine) of *E. coli* BL21(DE3) expressing SL52 following FPLC purification. **Expected size of SL52: 41.0 kDa**. Lanes (1) crude lysate, (2) Precision Plus Protein Dual Color Standards ladder (Bio-Rad), (3) pellet, (4) clarified lysate, (5) flow-through, (6) wash A, (7) wash 10% B, (8) fractions 5, 6, 7, 8, 10, 12, 14, 16. **E) SL226:** SDS-PAGE gels (Tris-glycine) of *E. coli* BL21(DE3) expressing SL226 following FPLC purification. **Expected size of SL226: 41.6 kDa**. Lanes (1) pre-solubilised pellet, (2) pellet, (3) clarified lysate, (4) flow-through, (5) wash A, (6) wash 10% B, (7) fractions 6, 7, 8, 10, 12, 14, 16, (8) Precision Plus Protein Dual Color Standards ladder (Bio-Rad). **F) SL235**: SDS-PAGE gels (Tris-glycine) of *E. coli* BL21(DE3) expressing SL235 following batch purification. **Expected size of SL235: 41.6 kDa**. Lanes (1) Precision Plus Protein Dual Color Standards ladder (Bio-Rad), (2) pre-solubilized pellet, (3) crude lysate, (4) pellet, (5) clarified lysate, (6) flow-through, (7) wash A, (8) wash 10% B, (9) wash 50% B-1, (10) wash 50% B-2, (11) wash 100% B-1, (12) wash 100% B-2. **G) SL237**: SDS-PAGE gels (Tris-glycine) of *E. coli* BL21(DE3) expressing SL237 following FPLC purification. **Expected size of SL237: 40.3 kDa**. Lanes (1) Precision Plus Protein Dual Color Standards ladder (Bio-Rad), (2) pre-solubilized pellet, (3) pellet, (4) clarified lysate, (5) flow-through, (6) wash A, (7) wash A-2, (8) wash 10% B, (8) fractions 6, 7, 8, 10, 12, 14, 16. **H) SL285**: SDS-PAGE gels (Tris-glycine) of *E. coli* BL21(DE3) expressing SL285 following batch purification. **Expected size of SL285: 41.5 kDa**. Lanes (1) Precision Plus Protein Dual Color Standards ladder (Bio-Rad), (2) pre-solubilized pellet, (3) crude lysate, (4) pellet, (5) clarified lysate, (6) flow-through 1, (7) flow-through 2, (8) wash A, (9) wash 10% B, (10) wash 50% B, (11) wash 100% B; **I)**: **SL306**: SDS-PAGE gels (Tris-glycine) of *E. coli* BL21(DE3) expressing SL306

following FPLC purification. **Expected size of SL306: 41.6 kDa**. Lanes (1) Precision Plus Protein Dual Color Standards ladder (Bio-Rad), (2) pre-solubilized pellet, (3) crude lysate, (4) pellet, (5) clarified lysate, (6) flow-through 1, (7) flow-through 2, (8) wash A, (9) wash 10% B, (10) wash 50% B, (11) wash 100% B.

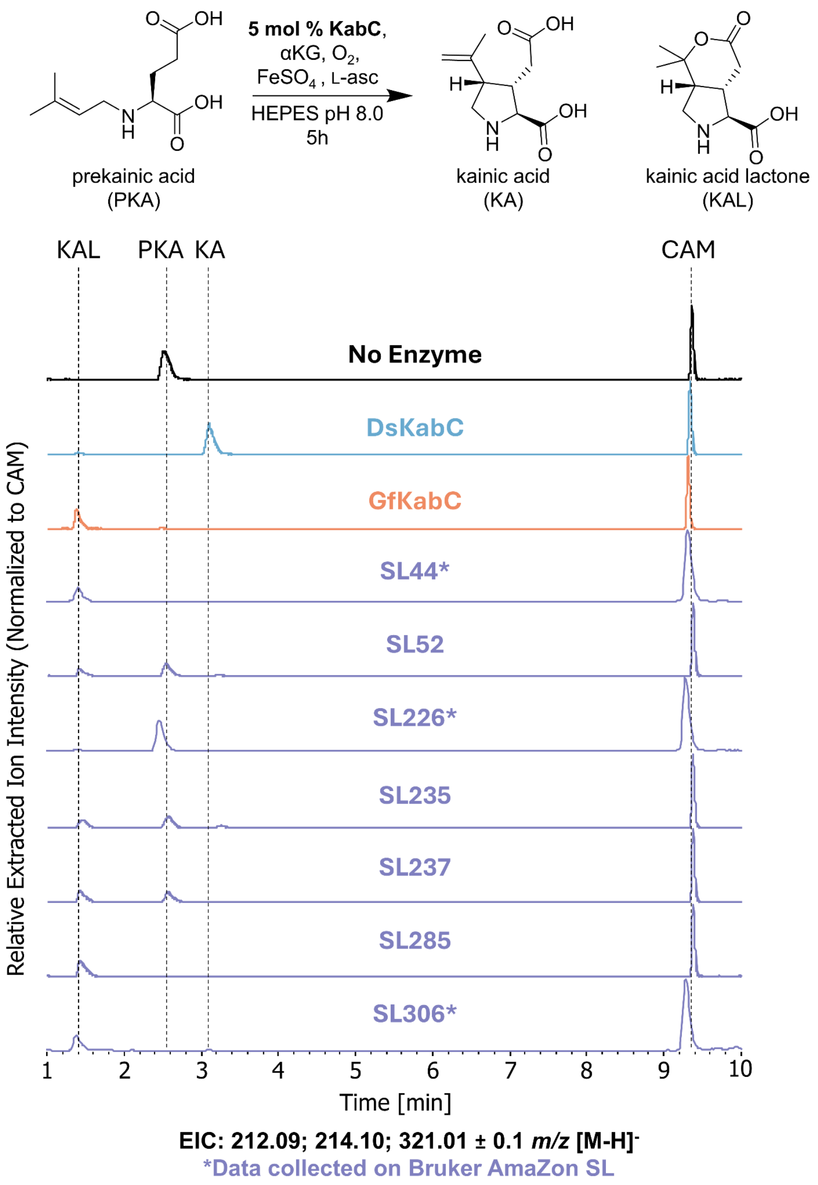


**Figure S18.** EICs of DsKabC, GfKabC, and all seven DNA shuffled variants selected for *in vitro* characterization with purified enzymes. CAM = 50mM chloramphenicol internal standard.

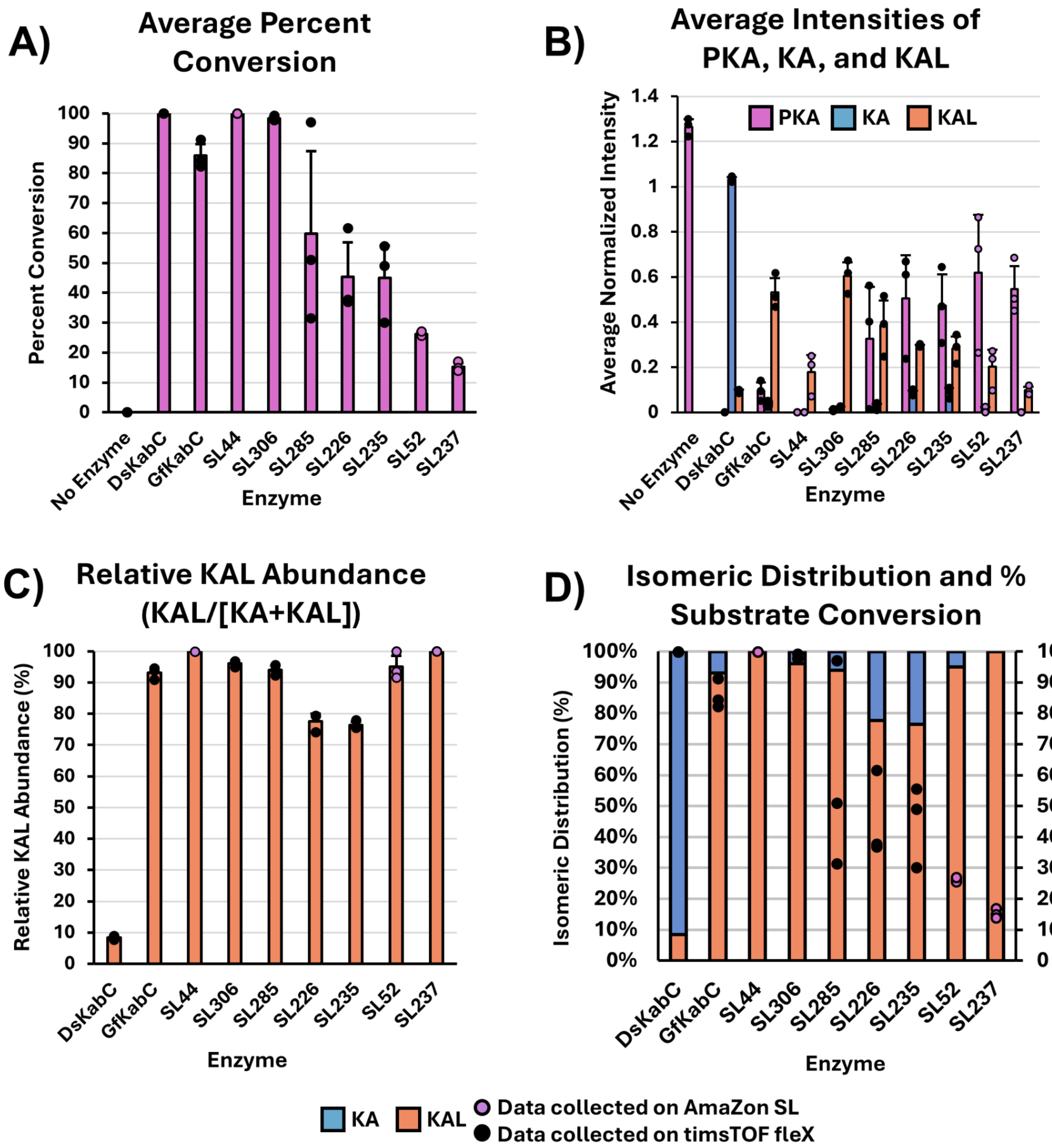


**Figure S19.** LC-MS Conversion data for KabC enzymes. A) Average percent conversion of PKA to kainoid products across 3 reaction replicates. B) Average intensities of PKA, KA, and KAL across 3 reaction replicates, normalized to chloramphenicol internal standard. C) Relative abundance of KAL across 3 reaction replicates. D) Average isomeric distribution (relative abundance) of KA and KAL with percent substrate conversion overlaid, across 3 reaction replicates.

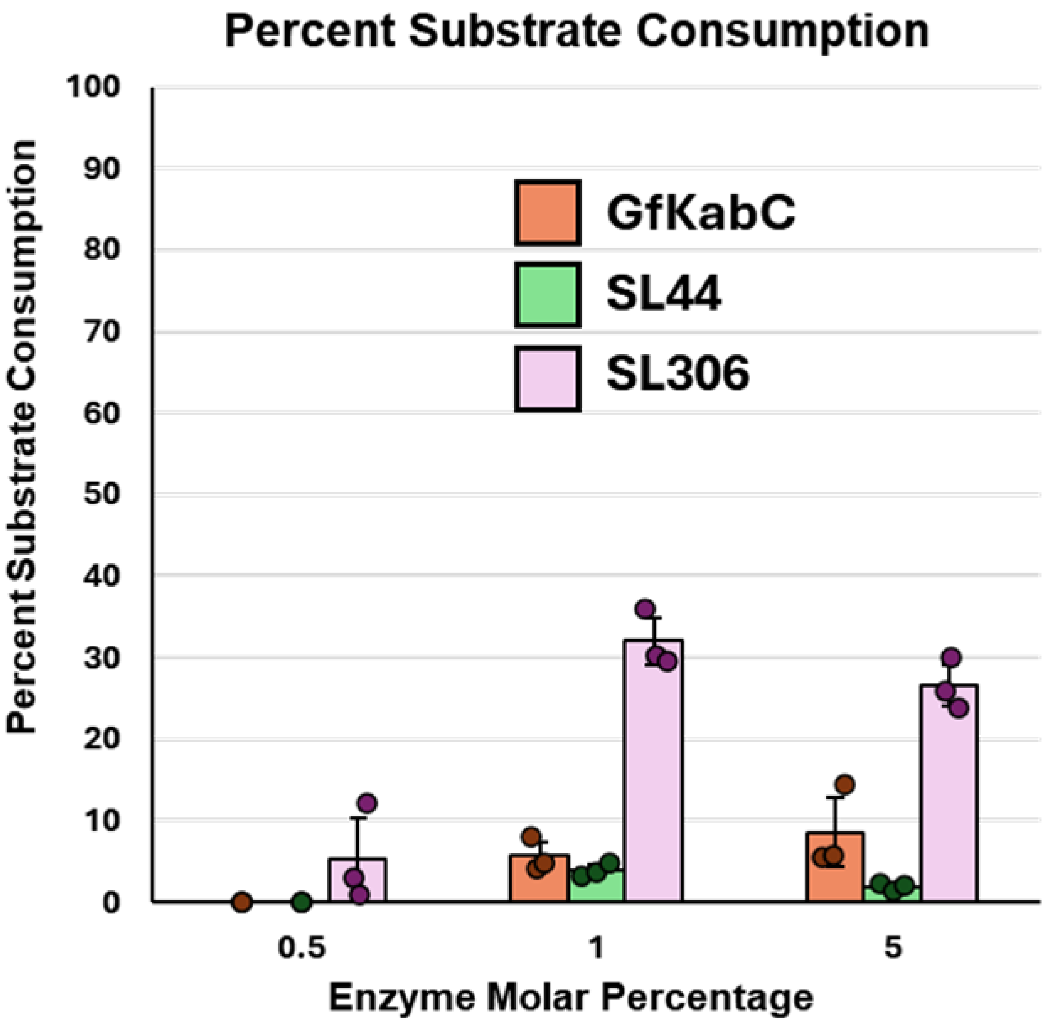


**Figure S20.** *In vitro* stability assessment of GfKabC, SL44, and SL306. *In vitro* activity assays for all three enzymes were set up as described in the Experimental Methods and Procedures (S47). Purified kainoid synthases were stored at -70°C for 4 months prior to thawing and testing again for *in vitro* activity. Enzyme concentrations were varied from 5 mol% (50 μM), down to 0.5 mol% (5 μM) and the reactions were carried out as previously described. The data displayed shows the substrate consumption of the PKA substrate across three reaction replicates, across three enzyme concentrations (Substrate consumption = [(1-(PKA/(PKA+KA+KAL)) * 100]). All intensities were normalized to a 50 μM chloramphenicol internal standard.

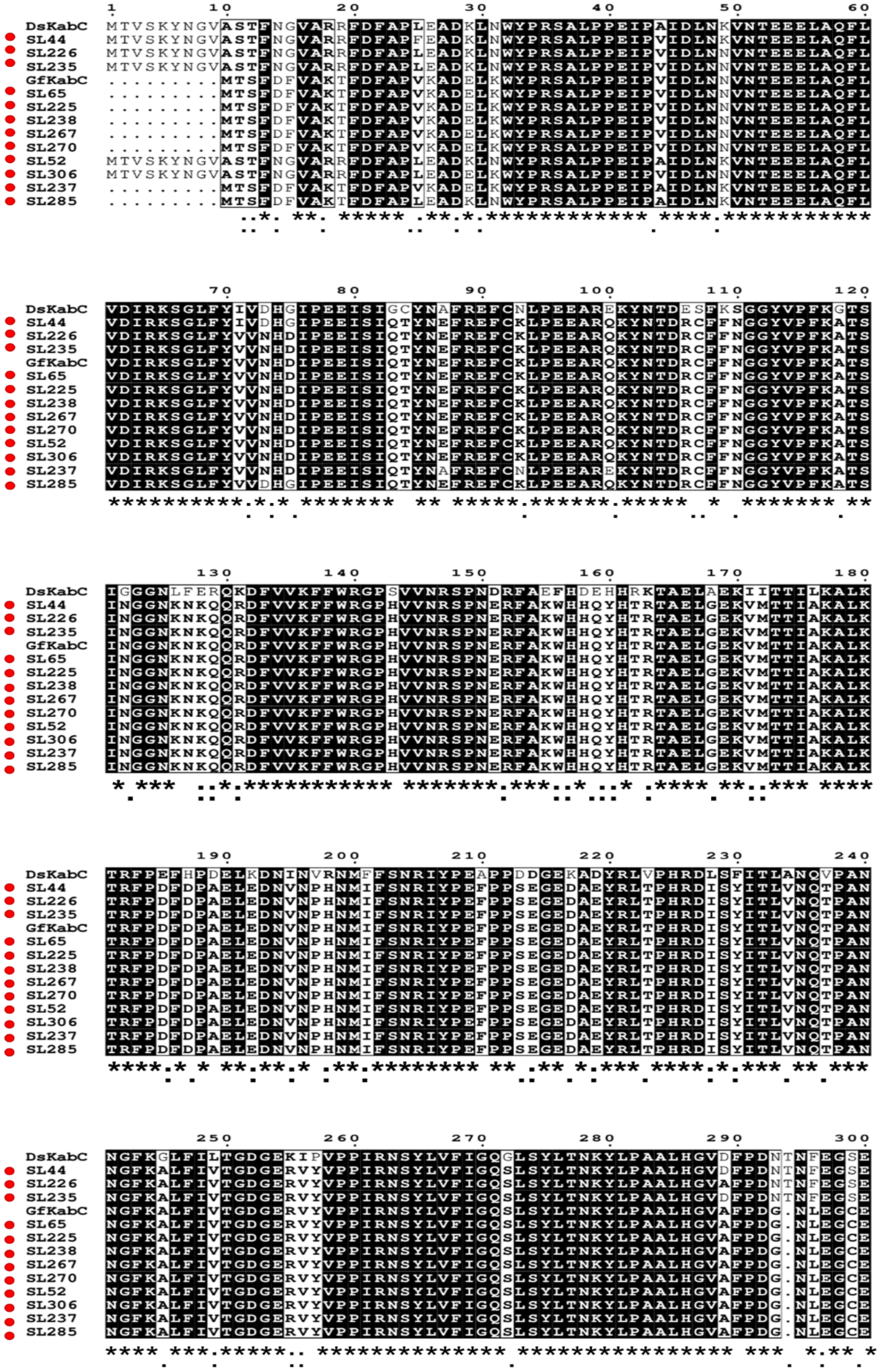
DsKabC
SL44
SL226
SL235
GfKabC
SL65
SL225
SL238
SL267
SL270
SL52
SL306
SL237
SL285
1
10
20
30
40
50
60
70
80
90
100
110
120
130
140
150
160
170
180
190
200
210
220
230
240
250
260
270
280
290
300

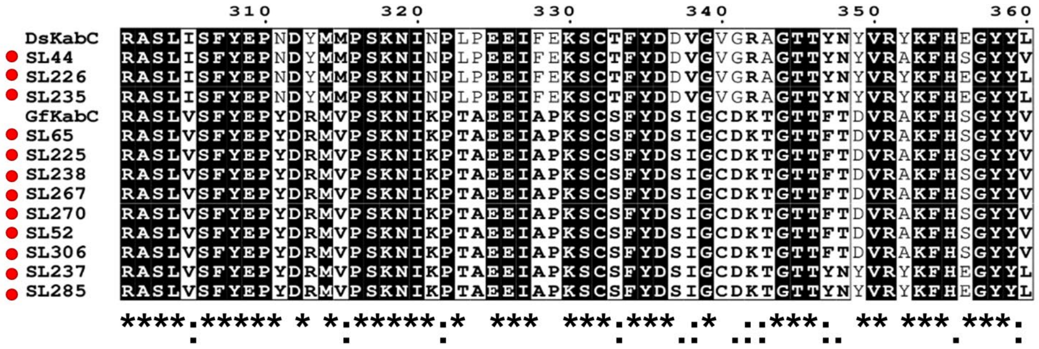

**Figure S21**: Multiple sequence alignment of the 12 hits that were sequenced with DsKabC and GfKabC using Clustal Omega[1] with visualisation performed using ESPript version 3.0.[2] Sequences SL65, SL225, SL238, SL267, and SL270 had the same sequence as that of GfKabC. 7 hits, i.e., SL44, SL52, SL226, SL235, SL237, SL285, and SL306 had shuffled sequences (denoted by red circles).

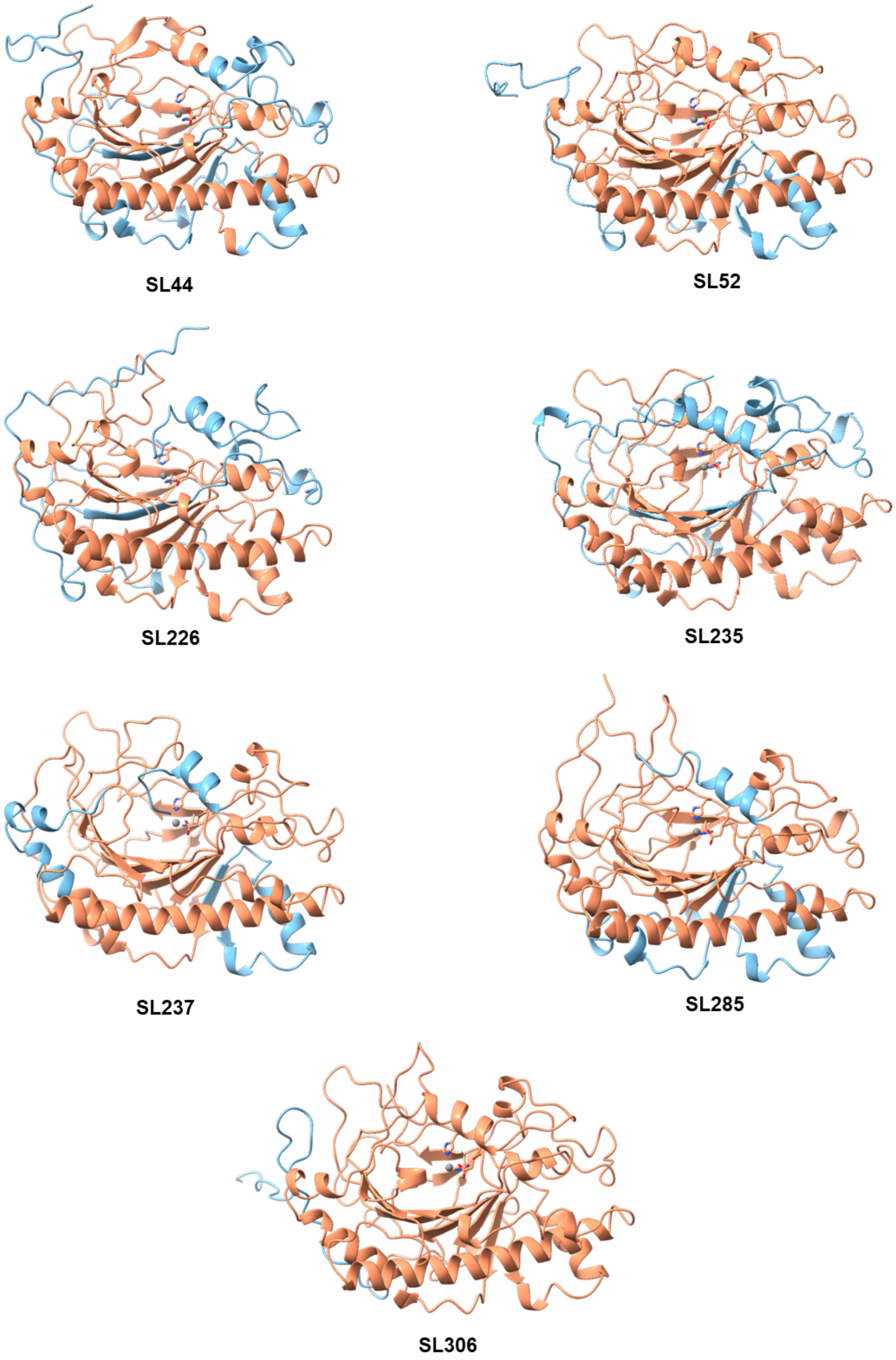


**Figure S22.** AlphaFold 3[3]-generated structure of the seven shuffle variants that are high KAL producers. Portions of the sequences that correspond to DsKabC and GfKabC are colored blue and orange respectively.

**Table S1. LB Media Recipe (Per 1L of Milli-Q $H_2O$) and preparation notes**

| Component | Amount (g) |
|---|---|
| Tryptone | 10 |
| Yeast extract | 5 |
| NaCl | 10 |
| Agar | 15 |
| pH 7.0 (adjust with 1M NaOH) | |
| **With kanamycin, prekainic acid, and/or IPTG*** | |
| 50 mg/mL kanamycin | 1 mL |
| 50mM PKA stock | 20 mL |
| 1M IPTG stock | 0.1 mL |

Autoclaved at 121°C for 20 minutes

*Stock solutions were sterile filtered and added post-sterilization once the media cooled to ~50°C.

- For standard LB-KAN agar plates, 20mL of agar media were transferred into a 100mm petri dish.
- For agar plates used for microbial MSI, 10mL of agar media were transferred into a 100mm petri dish. These plates are referred to as “thin agar”, and are used to ensure proper flattening of microbial material during the drying process for MALDI-MSI sample preparation.

**Table S2. Software Information**

| Software | Version |
|---|---|
| timsControl | 6.0.3_abe6ecc8_1 |
| flexImaging | 7.5 |
| SCiLS Lab | 2025b Pro |
| DataAnalysis | 6.1 (Build 213.27.0)(64-bit) |
| MALDI PharmaPulse | 2023c 3.2.0 build-2 |

**Table S3. MALDI PharmaPulse Campaign Parameters**

| ***m/z* Extraction Tolerance** | | | | | 20 ppm | | | |
|---|---|---|---|---|---|---|---|---|
| **Target List** | | | | | | | | |
| **Cmpd. #** | **Analyte** | **Compound ID** | **MF** | ***m/z*** | **Ion** | **1/K0 Start** | **1/K0 End** | **Response Factor** |
| 1 | Analyte | KA | - | 212.0893 | $[M\text{-}H]^-$ | 0.676 | 0.6925 | 1.0 |
| 2 | Analyte | KAL | - | 212.0894 | $[M\text{-}H]^-$ | 0.6925 | 0.71 | 1.0 |
| **Compound Ratio to Report** | | | | | | | | |
| **Numerator** | | | | | 212.0894 | | | |
| **Denominator** | | | | | 212.0893 | | | |
| **Monitoring** | | | | | | | | |
| Perform First level check: noise spectra | | | | | | | | |
| Max. number of consecutive noise spectra | | | | | 10 | | | |

**Note:** MALDI PharmaPulse will trigger a warning if the two *m/z* values of interest are exactly identical, as it is designed for isobaric analysis. To circumvent this warning, the user can offset the last decimal place by 0.0001 Da and still remain within the monitoring tolerance for each isomeric analyte. The software manufacturer was consulted on this approach for campaign design.

**Table S4. Instrument Parameters for MALDI-TIMS-MS**

| MS Settings | | $MS^1$ | $MS^2$ |
|---|---|---|---|
| Scan range | | 50-700 *m/z* | 50-700 *m/z* |
| Ion polarity | | negative | negative |
| Scan mode | | MS | prm-PASEF* |
| **Tune** | | | |
| Transfer | MALDI Plate Offset | 50.0 V | 50.0 V |
| | Deflection 1 Delta | -60 V | -60 V |
| | Funnel 1 RF | 250.0 Vpp | 250.0 Vpp |
| | isCID Energy | 0.0 eV | 0.0 eV |
| | Funnel 2 RF | 200.0 Vpp | 200.0 Vpp |
| | Multipole RF | 300.0 Vpp | 300.0 Vpp |
| Collision Cell | Collision Energy | 10.0 eV | 10.0 eV |
| | Collision RF | 400.0 Vpp | 400.0 Vpp |
| Quadrupole | Ion Energy | 5.0 eV | 5.0 eV |
| | Low Mass | 50.0 *m/z* | 50.0 *m/z* |
| Focus Pre TOF | Transfer Time | 50.0 µs | 50.0 µs |
| | Pre Pulse Storage | 3.5 µs | 3.5 µs |
| **Laser** | | | |
| Application | | MS Dried Droplet | MS Dried Droplet |
| Power Boost | | 0.0% | 0.0% |
| Scan Range | | X= 26.0 µm, Y= 26.0 µm | X= 26.0 µm, Y= 26.0 µm |

| | | |
|---|---|---|
| Resulting Field Size | X= 104.0 μm, Y= 104.0 μm | X= 104.0 μm, Y= 104.0 μm |
| Laser Power | 50% | 50% |
| Bursts | 5 | 5 |
| Shots | 1000 | 1000 |
| Frequency | 5000 Hz | 5000 Hz |
| **TIMS** | | |
| $1/K_0$ start | 0.67 | 0.67 |
| $1/K_0$ end | 0.73 | 0.73 |
| Ramp time | 200.0 ms | 200.0 ms |
| Accumulation time | 200.0 ms | 200.0 ms |
| Duty cycle | 100.0% | 100.0% |
| Ramp Rate | 4.85 Hz | 4.85 Hz |
| **Offsets** | | |
| Δt1 (deflection transfer ->capillary exit) | 20.0 V | -20.0 V |
| Δt2 (deflection transfer ->deflection discard) | 120.0 V | -120.0 V |
| Δt3 (funnel 1 in ->deflection transfer) | -15 V | -15V |
| Δt4 (accumulation trap ->funnel 1 in) | -50 V | -50 V |
| Δt5 (accumulation exit->accumulation transfer) | 0.0 V | 0.0 V |
| Δt6 (ramp start->accumulation exit) | -20.0 V | -20.0 V |

| | | |
|---|---|---|
| Collision cell in | -140.0 V | -140.0 V |
| **Advanced Parameters** | | |
| Lock accumulation to mobility range | Yes | Yes |

**Note:** Parameters are the same for MALDI-MS and MSI experiments, disregarding the TIMS parameters.

*PRM-PASEF Acquisition: *m/z* 212.09 was monitored from $1/K_0$= 0.68-0.72 $Vs/cm^2$

### Materials and Methods

#### Reagents

PKA, KA, and KAL were all prepared using previously established protocols.[4,5] Phosphorus red was purchased from Spectrum Chemical MFG Corporation (New Brunswick, NJ), and Agilent Tune Mix was purchased from Agilent Technologies Inc. (Santa Clara, CA).

#### Preparation of kainoid standard solutions for MS analysis

A small amount of KA, KAL, and PKA were suspended in 1:1 LC/MS-grade ACN and Milli-Q ultrapure $H_2O$ to form 1 mM stock solutions. The stocks were diluted 100-fold to form 0.01 mM (10 µM) solutions for analysis. For MALDI-TIMS-MS analysis, equal parts of the 10 µM KA and KAL solutions were mixed further suspended into an equal volume of 50 mM 1,5-DAN (in 1:1 ACN/$H_2O$). The matrix-analyte solution (1 µL) was spotted in duplicate onto a MALDI target plate and allowed to dry in a chemical fume hood before analysis on the timsTOF fleX mass spectrometer. The same 1:1 mixture of KA and KAL (without MALDI matrix) was directly infused into the timsTOF fleX and subjected to ESI-TIMS-MS analysis (Figure 2A).

#### MALDI-TIMS-MSI Sample Preparation

pET-28a(+) plasmid containing DsKabC was transformed into *E. coli* BL21 (DE3) competent cells, and the recovery culture was incubated at 37 °C until reaching an $OD_{600}$ of 0.1. A 5 µL aliquot of the recovery culture was then spotted in duplicate onto thin LB agar plates containing 50 µg/mL kanamycin, 1 mM PKA, and either with or without 0.1 mM isopropyl β-D-1-thiogalactopyranoside (IPTG). *E. coli* cultures were incubated inverted for 10 hrs at 37°C to promote colony formation. The plates were then transferred to a 30°C incubator for an additional 6 hrs.

Following incubation, the *E. coli* colonies along with the surrounding agar (~0.5 cm around the edge of each bacterial colony) were excised from the agar plates using a sterile razor blade and gently transferred to a ground steel MALDI target plate as outlined by Yang et al.[6] An optical image of the target plate with the colonies was taken prior to matrix application. A standard solution of 10 μM 1:1 KA:KAL was spotted on an empty spot for tuning the TIMS before acquisition. A 3:1 mixture of 1,5-diaminonaphthalene and 2,5-dihydroxybenzoic acid (3:1 1,5-DAN:2,5-DHB) was used as MALDI matrix and applied with a 53 micron stainless steel sieve (Hogentogler & Co, USA). The target plates were allowed to dry at 35°C for 4 hrs using a homemade spinning apparatus.[7] Following drying, excess matrix was removed using compressed air, and 1 μL of saturated red phosphorus in 1:1 ACN/$H_2O$ was spotted as a calibrant and allowed to dry. Finally, a second optical image of the target plate was taken following matrix application for subsequent MSI acquisition.

**Translation of biological sample preparation to colony-based format**

To test sample preparation, we performed transformations of DsKabC, *vide supra*. However, instead of spotting a small volume of recovery culture, we spread the culture evenly across the agar surface to promote colony formation across the entire plate, allowing for colony picking and direct transfer to MALDI targets. After incubation, single colonies were then transferred to spots on a MALDI target using sterile toothpicks, taking care to spread as uniformly as possible to minimize sample heterogeneity between spots. When screening mutant libraries, control spots should also be included for normalization. For library generation and storage, the residual material from each colony was inoculated into the corresponding well of a 96-deepwell plate containing LB broth with 50 μg/mL kanamycin. These 96-well culture plates were then incubated overnight at 37°C, and 50:50 glycerol-water was subsequently added to allow for storage at -70 °C until ready for sequencing. Once colony-transfer was complete, the colony material was allowed to dry on the surface of the MALDI target for ten min prior to matrix application. A 5 mg/mL solution of 3:1 1,5-DAN:2,5-DHB in 1:1 MilliQ $H_2O$:ACN was applied using a TM Sprayer (HTX Imaging). Matrix was sprayed rather than sieved or spotted to ensure the highest possible uniformity between spots, enabling reliable relative quantitation of KA and KAL. When assessing experimental reproducibility among 92 replicates of *E. coli* expressing DsKabC

Y347F, the coefficient of variation (CV) was ~17%, indicating variation adequate for first-pass screening (Figure S10).

**Instrumentation and Software**

MALDI-TIMS-MS data were collected on a timsTOF fleX (Bruker Daltonics) from 50-700 *m/z* and $1/K_0$ 0.67-0.73 $Vs/cm^2$ in negative ion mode. LC-MS data were collected either on the timsTOF fleX using the same parameters as for MALDI-MS, or on an AmaZon SL ion trap mass spectrometer (Bruker Daltonics) from *m/z* 100-2200, both equipped with an Elute UHPLC. An Intensity Solo 2 C18 column (100 x 2.1 mm; 1 µm; 100 Å; BRHSC18022100; Bruker Daltonics) was used for LC gradient analysis. LC analysis was performed with $H_2O$(A):ACN(B) w/ 0.1% formic acid using the following gradient: 0-3 min held at 95% A followed by a decrease over 3-6 min to 85% A with a further decrease over 6-12 min to 0% A and held from 12-14 min, followed by an increase over 14-15 min 95% A and finally reequilibrated from 15-17 min at 95% A. The flow rate of 0.3 mL/min. A full list of MS tune parameters can be found in the supplementary material (Table S4). Phosphorus red was used for mass calibration in MALDI mode, while Agilent Tune Mix was used for TIMS and/or LC-ESI-MS calibration. The software and versions used for all analyses is listed in Table S2. High-throughput data visualization was performed using a combination of MALDI-PharmaPulse and an in-house Python script. Semi-preparative HPLC for purifying PKA and KAL was performed using a 1260 Infinity (Agilent Technologies Inc.) equipped with a Synergi Polar-RP column (250 x 10mm; 4 µm; 80 Å; H20-415613; Phenomenex). The gradient used $H_2O$(A):ACN(B) w/ 0.1% formic acid in the following intervals: 0-5 min held at 95% A followed by a decrease over 5-16 min to 35% A, followed by an increase over 16-17 min to 95% A, decreased over 17-20 min to 5% A, then increased over 20-21 min to 95% A and held from 21-24 min at 95% A. The flow rate was 3.00 mL/min.

### MALDI matrix recrystallization

MALDI matrices α-cyano-4-hydroxycinnamic acid (CHCA) [(98%), Sigma-Aldrich] and 2,5-dihydroxybenzoic acid (2,5-DHB) [(98%), Sigma-Aldrich] were recrystallized in-house as previously described.[8] For recrystallization of 1,5-diaminonaphthalene (1,5-DAN) [D21200-100G, (98%), Sigma-Aldrich], 1500 mL of 90% ACN were transferred to a beaker with a magnetic stir bar. The solvent was then placed on a hot plate and heated to 70°C. A large watch glass was placed over the beaker to condense most of the solvent, but 100-150mL of 90% ACN was added during heating to compensate for evaporation. 100 g of 1,5-DAN were added to the solvent and the solution was heated up to 85°C, and stirred until the solution became transparent. The stir rate was then increased and the heat turned off, allowing for slow cooling overnight. The resulting crystals were filtered and collected using a Buchner funnel, and rinsed with cold 70% ACN. The crystals were dried overnight and the process was repeated once over to ensure the highest purity.

### High-throughput data visualization and variant selection

Data acquisition was performed using both timsControl and the MPP software, which operate in parallel. MPP provides an interface that is specifically designed to set up high-throughput screening experiments via MALDI-TIMS-MS. The user defines “substrates” and “analytes” in a target table (Table S3), designating the exact mass and the type of ion ($[M-H]^-$, $[M+Na]^+$, etc.) for each. With TIMS on, the user must also define the $1/K_0$ range for each analyte of interest. Once the target analyte table is filled out, the user can navigate to the “autoexport” tab and select from a list of columns that can be included in a *.csv file that is exported post-acquisition. The acquisition tab then allows the user to define spots for acquisition. Once the run is started, acquisition is performed in timsControl, but live MS and mobilogram traces can be visualized in MPP. Post-acquisition, the user can define multiple normalizations in MPP, and can even monitor for percent conversion. In our case however, we wanted to monitor the relative abundance of KAL and NOT percent conversion, which required normalization to DsKabC template control spots. This form of normalization cannot be performed directly in the MPP interface. For our purposes, we exported the generated *.csv file containing MALDI spot information, *m/z* values, and TIMS peak areas for each analyte for downstream analysis via a homemade python script. The script first calculates the relative abundance of KAL for all spots using the following convention:

$$Relative\ KAL\ abundance = \frac{A_{KAL}}{A_{KA} + A_{KAL}}$$

where $A_{KA}$ = the TIMS peak area of kainic acid and $A_{KAL}$ = the TIMS peak area of kainic acid lactone. The script then calculates the average relative abundance of the user-defined template enzyme control spots. The relative abundance of KAL for each variant spot is then divided by the average relative KAL abundance for the template controls to give a readout of fold change:

$$Fold\ change = \frac{\left(\frac{A_{KAL}}{A_{KA} + A_{KAL}}\right)_{Variant}}{\langle\frac{A_{KAL}}{A_{KA} + A_{KAL}}\rangle_{Template}}$$

where slanted brackets represent the average relative abundance of KAL for the template. The fold change values for each spot are visualized as an intensity heatmap organized by MALDI-spot number, with blue indicating a high fold change and white indicating a fold change of zero (or inactive/ no KAL production). Spots that do not meet a user-defined TIMS area threshold (for the total area of KA and KAL) are colored black on the heatmaps to minimize the selection of false positives or inactive variants (no conversion to products = no peak area to measure for KA or KAL). Lastly, variants with relative KAL abundance within three standard deviations of the template average are marked by a red X to identify functional mutants without improved relative

KAL production. It is important to reiterate that our selection criteria for this specific case are solely based on relative KAL abundance, which does NOT necessarily translate to improved substrate conversion. The script used for this analysis along with documentation can be found here: https://github.com/rashepherd/foldchangeviz. Lastly, development of an open-source command line tool for users without access to the MPP software is underway. This tool and its associated documentation can be found here: https://github.com/rashepherd/timsMAP. Please note that work on the timsMAP repository is ongoing and subject to community-based updates.

### Molecular Biology and Biochemical Methods

**General Considerations:** Chemically competent *E. coli* BL21(DE3) cells were prepared and used for cloning. Phusion HF Polymerase, 5X Phusion HF Buffer, DMSO, and dNTPs were purchased from New England Biolabs (NEB). QiAprep Spin Miniprep and Qiaquick PCR Purification Kits were purchased from Qiagen. The expression vector pEt28a(a)(+) was used from frozen glycerol stocks. All primers were purchased from IDT Technologies. A Bio-Rad T100™ Thermal Cycler and an Eppendorf Mastercycler were used for all PCRs.

#### Site Directed Mutagenesis of DsKabC

Primers containing the mutagenic codons were designed in-house and then synthesized (Integrated DNA Technologies). Then, site-directed mutagenesis PCR reactions were conducted using the wild-type *dsKabC* pET28a(+) plasmid as a template. After transformation into *E. coli* DH106 cells, plasmid purification, and confirmation of successful mutants via Sanger sequencing (Azenta Life Sciences), the mutagenized plasmids were transformed into *E. coli* BL21(DE3) cells for heterologous protein expression following the same protocols discussed below.

#### Construction of KabC Fusion Enzymes

Each of the fusion genes was synthesized and cloned into pET28a(+) vectors with *N*-terminal $His_6$ tags (Twist Biosciences). Fusion plasmids were then transformed and expressed in BL21(DE3) cells following the protocols discussed below. Initial whole cell lysate assays were then conducted to assess whether these fusion proteins had any apparent change in their KA:KAL product ratios compared to the wild-type DsKabC (Figure S5).

**Codon-optimized DNA and Protein Sequences**

***dsKabC* from *Diginea simplex***

ATGACAGTAAGTAAGTACAACGGTGTTGCGAGTACGTTCAACGGTGTTGCAAGGAGATTCG
ACTTTGCCCCATTGGAGGCAGATAAATTGAATTGGTATCCCAGGTCAGCTCTTCCTCCAGA
AATCCCTGCTATCGACCTGAATAAGGTCAATACTGAAGAAGAACTGGCTCAATTCCTGGTC
GATATTCGCAAATCTGGGCTCTTTTATATTGTTGATCATGGCATTCCAGAGGAGATCTCTAT
AGGATGTTACAATGCATTCCGTGAGTTTTGCAACCTTCCCGAGGAAGCAAGAGAGAAGTAC
AACACAGATGAGTCCTTTAAAAGCGGTGGCTATGTTCCTTTCAAAGGCACGTCGATTGGTG
GAGGGAACTTGTTTGAACGACAGAAGGATTTCGTTGTCAAATTCTTTTGGAGAGGACCAAG
TGTTGTCAATAGGTCTCCTAATGATCGCTTTGCCGAATTCCATGATGAACATCATCGAAAGA
CAGCTGAGTTGGCCGAAAAAATCATCACTACCATTTTGAAAGCCCTGAAGACACGCTTTCC
AGAATTCCATCCTGACGAGCTAAAGGATAATATCAATGTTCGAAACATGTTTTTCAGTAATC
GAATCTATCCAGAGGCTCCGCCAGATGATGGAGAAAAAGCTGACTACCGACTTGTTCCTCA
TCGAGATCTCAGTTTTATCACTCTTGCAAATCAAGTTCCGGCTAACAATGGATTCAAGGGTC
TTTTTATACTGACAGGTGATGGAGAAAAAATTCCTGTTCCTCCAATCCGGAACAGCTACTTG
GTCTTCATCGGTCAAGGCCTCTCATATCTCACGAATAAGTATCTTCCTGCAGCGCTTCATG
GTGTGGACTTTCCAGACAATACAAATTTTGAAGGAAGTGAAAGAGCCTCTCTGATCAGCTT
CTACGAGCCAAACGATTACATGATGCCATCCAAGAACATTAACCCTTTACCTGAGGAGATA
TTTGAGAAAAGCTGTACGTTTTACGATGATGTCGGAGTAGGGAGGGCTGGTACAACATATA
ATTACGTGAGGTACAAATTTCATGAAGGATACTACCTTTAA

**DsKabC Protein Sequence (accession number QCC62383.1)**

MTVSKYNGVASTFNGVARRFDFAPLEADKLNWYPRSALPPEIPAIDLNKVNTEEELAQFLVDIR
KSGLFYIVDHGIPEEISIGCYNAFREFCNLPEEAREKYNTDESFKSGGYVPFKGTSIGGGNLFER
QKDFVVKFFWRGPSVVNRSPNDRFAEFHDEHHRKTAELAEKIITTILKALKTRFPEFHPDELKD
NINVRNMFFSNRIYPEAPPDDGEKADYRLVPHRDLSFITLANQVPANNGFKGLFILTGDGEKIPV
PPIRNSYLVFIGQGLSYLTNKYLPAALHGVDFPDNTNFEGSERASLISFYEPNDYMMPSKNINPL
PEEIFEKSCTFYDDVGVGRAGTTYNYVRYKFHEGYYL

***gfKabC* from *Grateloupia filicina*** (*E. coli* codon-optimized and optimized to match *dsKabC*)

ATGACTTCTTTCGATTTCGTTGCAAAGACTTTCGACTTTGCCCCAGTCAAAGCAGATGAGTT
GAAGTGGTATCCCAGGTCAGCTCTTCCTCCAGAAATCCCTGTAATCGACCTGAATAACGTC
AATACTGAAGAAGAACTGGCTCAATTCCTGGTCGATATTCGCAAATCTGGGCTCTTTTATGT
TGTTAACCATGATATTCCAGAGGAGATCTCTATACAGACGTACAATGAATTCCGTGAGTTTT
GCAAACTTCCCGAGGAAGCAAGACAAAAGTACAACACAGATCGCTGCTTTTTCAATGGTGG
CTATGTTCCTTTCAAAGCCACGTCGATTAATGGAGGGAACAAGAACAAACAGCAGCGCGAT
TTCGTTGTCAAATTCTTTTGGAGAGGACCACACGTTGTCAATAGGTCTCCTAATGAACGCTT
TGCCAAATGGCATCATCAGTATCATACACGCACAGCTGAGTTGGGTGAAAAAGTCATGACT
ACCATTGCAAAAGCCCTGAAGACACGCTTTCCAGACTTCGACCCTGCCGAGCTAGAGGAT
AATGTGAATCCTCACAACATGATTTTCAGTAATCGAATCTATCCAGAGTTCCCGCCAAGTGA
AGGAGAAGATGCTGAGTACCGACTTACCCCTCATCGAGATATTAGTTACATCACTCTTGTG
AATCAAACTCCGGCTAACAATGGATTCAAGGCACTTTTTATAGTAACAGGTGATGGAGAAC
GCGTTTACGTTCCTCCAATCCGGAACAGCTACTTGGTCTTCATCGGTCAAAGCCTCTCATA
TCTCACGAATAAGTATCTTCCTGCAGCGCTTCATGGTGTGGCTTTTCCAGACGGCAATCTT
GAAGGATGCGAAAGAGCCTCTCTGGTGAGCTTCTACGAGCCATATGATCGCATGGTCCCA
TCCAAGAACATTAAACCTACGGCAGAGGAGATAGCTCCAAAAAGCTGTTCCTTTTACGATT
CGATTGGATGTGACAAGACAGGTACAACATTTACTGATGTGAGGGCAAAATTTCATTCGGG
ATACTACGTCTAA

**GfKabC Protein Sequence (accession number QCC62378.1)**

MTSFDFVAKTFDFAPVKADELKWYPRSALPPEIPVIDLNNVNTEEELAQFLVDIRKSGLFYVVNH
DIPEEISIQTYNEFREFCKLPEEARQKYNTDRCFFNGGYVPFKATSINGGNKNKQQRDFVVKFF
WRGPHVVNRSPNERFAKWHHQYHTRTAELGEKVMTTIAKALKTRFPDFDPAELEDNVNPHNM
IFSNRIYPEFPPSEGEDAEYRLTPHRDISYITLVNQTPANNGFKALFIVTGDGERVYVPPIRNSYL
VFIGQSLSYLTNKYLPAALHGVAFPDGNLEGCERASLVSFYEPYDRMVPSKNIKPTAEEIAPKSC
SFYDSIGCDKTGTTFTDVRAKFHSGYYV

**Error Prone PCR for Random KabC Mutagenesis**

Error-prone PCR (epPCR) was conducted using the Genemorph II random mutagenesis kit (Agilent), or by varying $MnCl_2$ concentrations based on previously established protocols.[9] DsKabC Y347F was used as the template given that this mutation already displayed increased relative production of KAL *in vitro*. The resulting PCR product was digested with restriction endonuclease dpnI to degrade the remaining parent plasmid and cloned into pET28a vector via Gibson Assembly, the product of which was then transformed into *E. coli* BL21(DE3) cells and plated onto LB+KAN agar plates containing 1 mM PKA. Individual colonies containing unknown DsKabC Y347F variants were transferred to a MALDI plate and subjected to MALDI-TIMS-MS analysis (Figure S12).

**DNA shuffling for chimeric KabC generation**

The procedure for DNA shuffling was adapted and modified from previously published protocols.[10–13]

**Preparing *dsKabC* and *gfKabC* linear DNA for shuffling:** The *dsKabC* and *gfKabC* genes were PCR-amplified from their respective parental pET28a(+) plasmid DNA in 50 µL reactions consisting of the components listed in Table S6 using the protocol listed below. The reactions were subjected to DpnI digestion to remove unreacted template DNA and were further purified using Qiagen's PCR Clean-up kit.

Initial denaturation at 98 °C (30 s), 25 cycles of 98 °C (10 s), 67 °C (30 s), 72 °C (2 min), a final elongation of 72 °C (5 min) and hold at 4 °C until removal of products from PCR thermocycler.

**DNase I Digestion to obtain fragments for shuffling:** 5 µL of 10X DNase I buffer (1 M Tris-HCl, pH 7.4, 200 mM $MnCl_2$) was added to a tube containing 1 µg of each PCR-amplified insert and incubated at 15 ºC for 5 minutes before adding 0.5 µL 1 U/µL DNase I. The mixture was digested at 15 ºC for 2 minutes and the reaction was terminated by incubation at 90 ºC for 10 minutes. The reaction was purified by performing a PCR Clean-up.

**Reassembly of DNA fragments**: Spin column-purified fragments were added to 10 µL 5X Phusion HF buffer, 1 µL of 10 mM dNTP mix, 0.5 µL of Phusion DNA Polymerase and assembled using the following program:

Initial denaturation at 96 °C, 90 s; 35 cycles of (94 °C, 30 s; 65 °C, 90 s; 62 °C, 90 s; 59 °C, 90 s; 56 °C, 90 s; 53 °C, 90 s; 50 °C, 90 s; 47 °C, 90 s; 44 °C, 90 s; 41 °C, 90 s; 72 °C, 4 min); a final elongation at 72 °C, 7 min; and hold at 4 °C.

**PCR Amplification of reassembled DNA**: 1 µL of a 1:10 dilution of the unpurified reassembly reaction was subjected to PCR amplification in 100 µL reactions consisting of the components listed in Table S7 using the protocol listed below to obtain two distinct bands corresponding to . the sizes of *dsKabC* (1083 bp) and *gfKabC* (1053 bp). The corresponding bands were excised and purified by agarose gel purification.

**Cloning of the shuffled mutants**:

The reassembled DNA was sub-cloned into the pET28a(+) kanamycin resistant expression vector containing an N-terminal hexahistidine ($His_6$) tag using the primers listed in Table S5. Linearized pET28a(+) backbone was prepared by PCR amplification in a 250 µL reaction consisting of the components listed in Table S8 using the protocol listed below.

Initial denaturation at 98 °C (2 min), 30 cycles of 98 °C (10 s), 64.4 °C (30 s), 72 °C (2:30 min), a final elongation of 72 °C (2 min) and hold at 4 °C until removal of products from PCR thermocycler.

A Gibson assembly[14] was performed with the HiFi Assembly polymerase mix (New England Biolabs Inc.) and incubated at 55 ºC for 60 minutes and transformed into *E. coli* BL21(DE3) cells.

**Transformation into *E. coli***

All plasmids in this work were transformed into *E. coli* BL21(DE3) cells for plasmid expression by heat shock. After thawing cell aliquots from -70 °C storage on ice for 10 minutes, 10 µL of the Gibson assembly reaction was added to cell aliquots and incubated on ice for 30 minutes. Cells were heat shocked at 42 °C for 30 seconds, then incubated on ice again for 2 minutes. Aliquots were added to 950 µL of LB media and then incubated at 37 °C at 200 rpm for 1 hour. The negative control plasmid (empty pET28a(+)) and the positive control plasmid (pET28a(+)-*dsKabC*) were simultaneously transformed into *E. coli* BL21(DE3). Afterward, cells were plated on LB media plates with 50 µg/mL kanamycin and 50 mM PKA and incubated at 37 °C for 16 hours followed by incubation at 30 ºC for 6 hours

**Plasmid Purification**

Plasmid purification was achieved using the QIAprep Spin Miniprep Kit (QIAGEN) with the manufacturer's protocol. DNA concentrations were measured via NanoDrop UV-vis spectrophotometry and sent for sequencing (Azenta Life Sciences).

**General heterologous protein expression in *E. coli* BL21(DE3)**

The expression and purification procedure was adapted from previously published protocols.[5] Either a colony of transformed BL21(DE3) cells or a glycerol stock was used to inoculate 30 mL of LB media supplemented with kanamycin (50 µM). The culture was grown overnight at 37 °C, 200 rpm. 10 mL of this overnight culture was added per liter of Terrific Broth supplemented with kanamycin (50 µM). Each liter of culture was grown in a shaking incubator at 37 °C, 200 rpm until OD600 reached ~0.6. The flasks were then cooled to 18 °C over 45 - 60 min at which point 0.5 mM IPTG was added to induce protein expression. The cultures were allowed to grow at 18 °C and 200 rpm overnight (16 - 18 h) and pelleted in a centrifuge at approx. 3500 x g for 30 min. Supernatant was discarded and the cell pellet was resuspended in 25 mL of lysis buffer (1 M NaCl, 20 mM Tris-HCl pH 8.0, 30 mM imidazole, and 10% glycerol) and stored at -70 °C. If cells were to be immediately purified, the pellet was resuspended in buffer A (1 M NaCl, 20 mM Tris- HCl pH 8.0, 30 mM imidazole) and put on ice.

**Ni-NTA FPLC affinity purification**

Resuspended cell pellets were lysed on ice via sonication at 40% amplitude alternating between 15 sec on and 45 sec off (5 minutes total 'on' time). The lysate was pelleted via centrifugation at 4 °C and ~16,000 x g for 60 minutes. The clarified lysate was loaded onto a 5 mL HisTrap FF column (GE Healthcare Life Sciences) that was pre-equilibrated with at least 5 CV of buffer A. After loading, the column was washed with buffer A until the UV absorbance plateaued and reached approximately 50 mAU. The column was then washed with 10% buffer B for 5 - 10 CV to remove any loosely bound proteins. The protein was then eluted using a linear gradient of 100% buffer A to 100% buffer B (1 M NaCl, 20 mM Tris-HCl pH 8.0, 250 mM imidazole) over 60 mL while collecting 5 mL fractions. The flowrate through the column was 2 mL/min throughout the purification process. Fractions were assessed for purity using SDS-PAGE. Fractions containing purified protein had 2 mM EDTA added to remove any bound metals and were combined and further purified through a PD-10 desalting column (GE Healthcare Life Sciences) pre-equilibrated with GF buffer (20 mM HEPES pH 8.0, 300 mM KCl) following the default gravity filtration protocol. The protein was concentrated by Amicon ultra centrifugal filters (30 kDa molecular weight cut-off). Enzyme concentration was determined via Bradford assay and pure enzyme was either used immediately for assays or aliquoted and stored at -70 °C.

**Batch purification of proteins**

Following the sonication and centrifugation protocol mentioned above, the clarified lysate was loaded on to a 5 mL Ni-NTA resin packed in a 5 mL column that was pre-equilibrated with at least 5 CV of Buffer A. After loading, the column was washed with 5 CV of Buffer A followed by 5 CV of 10% Buffer B to remove any loosely bound proteins. The protein was then eluted using 10 CV of 50% Buffer B and 10 CV of 100% Buffer B. Fractions were assessed for purity using SDS-PAGE. Fractions containing purified protein had 2 mM EDTA added to remove any bound metals and were combined and further purified through a PD-10 desalting column (GE Healthcare Life Sciences) pre-equilibrated with GF buffer (20 mM HEPES pH 8.0, 300 mM KCl) following the default gravity filtration protocol. The protein was concentrated by Amicon ultra centrifugal filters (30 kDa molecular weight cut-off). Enzyme concentration was determined via Bradford assay and pure enzyme was either used immediately for assays or aliquoted and stored at -70 °C.

**Analytical Shuffle Enzyme Assays with PKA**

Analytical enzyme assays were conducted in sterile filtered milliQ $H_2O$ with 1 mM L-ascorbate, 5 mM α-ketoglutarate, and 125 μM $FeSO_4$ with 1 mM of PKA and 25 μM of purified enzymes. The enzymes stored in GF buffer were diluted to 25 µM. Enzyme followed by $FeSO_4$ were the last additions to assay and the reaction was allowed to run overnight for 18 h. The total volume for the reaction was 100 µL. The reaction was then quenched with 0.1 mM chloramphenicol in methanol and centrifuged at ~ 21,000 x g for 5 min. The supernatant was then subjected to analysis by UPLC-MS.

**Table S5. Primers used in this study.** In the primers for DsKabC site-directed mutagenesis PCR, red indicates the mutagenic codon.

| | |
|---|---|
| Pet28_dsKabC_S107C_F | CACAGATGAGtgcTTTAAAAGC |
| Pet28_dsKabC_S107C_R | TTGTACTTCTCTCTTGCTTCC |
| Pet28_dsKabC_G122N_F | CACGTCGATTaatGGAGGGAACT |
| Pet28_dsKabC_G122N_R | CCTTTGAAAGGAACATAGCCACC |
| Pet28_dsKabC_L126K_F | TGGAGGGAACaagTTTGAACGA |
| Pet28_dsKabC_L126K_R | CCAATCGACGTGCCTTTGAAAG |
| Pet28_dsKabC_E128K_F | GAACTTGTTTaaaCGACAGAAGGATTTCG |
| Pet28_dsKabC_E128K_R | CCTCCACCAATCGACGTG |
| Pet28_dsKabC_ V197P_F | TAATATCAATcctCGAAACATGTTTTTCA |
| Pet28_dsKabC_V197P_R | TCCTTTAGCTCGTCAGGATG |
| Pet28_dsKabC_F201I_F | TCGAAACATGattTTCAGTAAT |
| Pet28_dsKabC_F201I_R | ACATTGATATTATCCTTTAGCTC |
| Pet28_dsKabC_V223T_F | CTACCGACTTactCCTCATCGAG |
| Pet28_dsKabC_V223T_R | TCAGCTTTTTCTCCATCATCTGG |

| | |
|---|---|
| Pet28_dsKabC_Y347F_F | TGGTACAACAtttAATTACGTGAGGTACAAATTTCA |
| Pet28_dsKabC_Y347F_R | GCCCTCCCTACTCCGACA |
| Outer_dsKabC_gfKabC_F | GTGAGCGGATAACAATTCCCCTCTAG |
| Outer_dsKabC_gfKabC_R | GGTTATGCTAGTTATTGCTCAGCGGTGG |
| Inner_dsKabC_F | GGTGCCGCGCGGCAGCCATATGACAGTAAGTAAGTACAACGGTGTTG |
| Inner_dsKabC_R | GGTGGTGGTGGTGCTCGAGTTAAAGGTAGTATCCTTCATGAAATTTG |
| Inner_gfKabC_wob_F | CTGGTGCCGCGCGGCAGCCATATGACTTCTTTCGATTTCGTTG |
| Inner_gfKabC_wob_R | GGTGGTGGTGGTGCTCGAGTTAGACGTAGTATCCCGAATG |
| Pet28_NdeI_F | CATATGGCTGCCGCGCGGCAC |
| Pet28_XhoI_R | CTCGAGCACCACCACCACCACCACTGA |

**Table S6.** Components of the PCR performed to obtain linear DNA for shuffling.

| **Component** | **Volume Added (uL)** | **Final Concentration** |
|---|---|---|
| 5X Phusion HF Buffer | 10 | 1X |
| 10 mM dNTPs | 1 | 200 μM |
| 10 μM Outer_dsKabC_gfKabC_F | 2.5 | 0.5 μM |
| 10 μM Outer_dsKabC_gfKabC_ R | 2.5 | 0.5 μM |
| pET28a(+)-*dsKabC* DNA / pET28a(+)-*gfKabC* DNA | 1.2 | 20 ng |
| DMSO | 1.5 | 3% |
| Phusion DNA Polymerase | 0.5 | 1.0 unit |
| Sterile milliQ H2O | 30.8 | N/A |
| | Total = 50 μL | |

**Table S7.** Components of the PCR performed to reamplify the reassembled DNA fragments for shuffling.

| **Component** | **Volume Added (uL)** | **Final Concentration** |
|---|---|---|
| 5X Phusion HF Buffer | 20 | 1X |
| 10 mM dNTPs | 2 | 200 µM |
| 10 µM Inner_dsKabC_F | 5 | 0.5 µM |
| 10 µM Inner_dsKabC_ R | 5 | 0.5 µM |
| 10 µM Inner_gfKabC_wob_F | 5 | 0.5 µM |
| 10 µM Inner_gfKabC_wob_R | 5 | 0.5 µM |
| Reassembled DNA | 1 | 8.5 ng |
| DMSO | 3 | 3% |
| Phusion DNA Polymerase | 1 | 1.0 unit |
| Sterile milliQ H2O | 53 | N/A |
| | Total = 100 µL | |

**Table S8.** Components of the PCR performed to obtain linearized pET28a(+) DNA.

| **Component** | **Volume Added (uL)** | **Final Concentration** |
|---|---|---|
| 2X Primestar Mix | 125 | 1X |
| 10 μM Pet28a_NdeI_F | 5 | 0.5 μM |
| 10 μM Pet28a_XhoI_R | 5 | 0.5 μM |
| pET28a(+)DNA | 1 | 1 ng |
| DMSO | 7.5 | 3% |
| Sterile milliQ H2O | 106.5 | N/A |
| | Total = 250 μL | |
